# Isolation and genomic and physiological characterization of *Parageobacillus* sp. G301, the isolate capable of both hydrogenogenic and aerobic carbon monoxide oxidation

**DOI:** 10.1101/2023.01.17.524042

**Authors:** Yoshinari Imaura, Shunsuke Okamoto, Taiki Hino, Yusuke Ogami, Yuka Adachi Katayama, Ayumi Tanimura, Masao Inoue, Ryoma Kamikawa, Takashi Yoshida, Yoshihiko Sako

## Abstract

Prokaryotes, known as carbon monoxide (CO) oxidizers, use CO as the carbon or energy source with CO dehydrogenases (CODHs), which are divided into nickel-containing CODH (Ni-CODH) that are sensitive to O_2_ and molybdenum-containing CODH (Mo-CODH) that are capable of aerobic functioning. The oxygen conditions for CO oxidizers to oxidize CO may be limited because CO oxidizers isolated and characterized so far have either Ni- or Mo-CODH. Here, we report a novel CO oxidizer capable of CO oxidation with both types of CODH based on genomic and physiological characterization of the isolate *Parageobacillus* sp. G301. This thermophilic facultative anaerobic Bacillota bacterium was isolated from the sediment of a freshwater lake. Genomic analyses showed that G301 was the only isolate possessing both Ni-CODH and Mo-CODH. Genome-based reconstruction of the respiratory machinery and physiological investigation indicated that CO oxidation by Ni-CODH was coupled with H_2_ production (proton reduction), and CO oxidation by Mo-CODH was coupled with O_2_ reduction under aerobic conditions and nitrate reduction under anaerobic conditions. G301 would thus be able to thrive via CO oxidation under a wide range of conditions, from aerobic environments to anaerobic environments even without terminal electron acceptors other than protons. As comparative genome analyses revealed no significant differences in genome structures and encoded cellular functions, except for CO oxidation between CO oxidizers and non-CO oxidizers in the genus *Parageobacillus*, CO oxidation genes would be retained exclusively for CO metabolism and related respiration.

**Importance:** Microbial CO oxidation has received a lot of attention because it contributes to global carbon cycling in addition to functioning as a remover of CO, which is toxic to many organisms. Microbial CO oxidizers have a punctate phylogenetic distribution throughout bacteria and archaea, even in genus-level monophyletic groups. In this study, we demonstrated that the new isolate *Parageobacillus* sp. G301 is capable of both anaerobic (hydrogenogenic) and aerobic CO oxidation, which had not been previously reported. The discovery of this new isolate, which is versatile in CO metabolism, would accelerate research into such CO oxidizers with diverse CO metabolisms, expanding our understanding of microbial diversity. Through comparative genomic analyses, we propose that CO oxidation genes are optional but not essential genetic elements in the genus *Parageobacillus*, providing insight into a factor that shapes the mosaic phylogenetic distribution of CO oxidizers, even in genus-level monophyletic groups.

## Introduction

Carbon monoxide (CO) is toxic to many organisms, including humans, owing to its high affinity for transition metals in metalloproteins, such as heme (1, 2). It is a thermodynamically favorable electron donor with an *E^o^*’ of -520 mV for the CO/carbon dioxide (CO_2_) redox pair (3, 4). As a result, CO oxidation can be coupled with the reduction of various terminal electron acceptors, such as O_2_, Fe(III), nitrate, perchlorate, sulfur compounds, CO_2_, and protons (5–9).

Some prokaryotes, known as CO oxidizers, couple CO oxidation with the reduction of a terminal electron acceptor to gain energy (6, 10–12). In addition to contributing to the carbon cycle by consuming the CO contained as a trace gas in the atmosphere (6), CO oxidizers would provide a “safety valve” to the surrounding microbial community by consuming toxic levels of CO (13). The key enzyme in CO oxidation is carbon monoxide dehydrogenase (CODH), which catalyzes the reversible conversion between CO and CO_2_. There are two distinct CODHs: nickel-containing CODH (Ni-CODH) (14) and molybdenum-containing CODH (Mo-CODH) (15).

The catalytic subunit of Ni-CODH, which belongs to the hybrid-cluster protein family (Pfam ID: PF03063) (16, 17), is divided into two types: bacterial CooS-type Ni-CODH and archaeal CdhA-type Ni-CODH. An earlier phylogenetic classification of Ni-CODH included six distinct clades: Clade A corresponds to CdhA-type Ni-CODH, while clades B, C, D, E, and F corresponds to CooS-type Ni-CODH (18). A recent categorization based on phylogenetic relationships and the structure of active site motifs identified two additional clades: clade G and clade H (14, 19). Enzymatic activity has only been biochemically or physiologically confirmed for four of the seven Ni-CODH clades, namely Ni-CODHs from clades A (20), C (21), E (22), and F (23). Although homologs of Ni-CODHs belonging to other clades have been found in diverse prokaryotic genomes (14), their catalytic activities and physiological roles remain to be characterized. CO oxidizers containing *cooS/cdhA* can couple CO oxidation with the reduction of one or more terminal electron acceptors, including protons, CO_2_, fumarate, sulfate, nitrate, and Fe(III) (3, 10, 24). One of the most well-known of these is hydrogenogenic CO oxidizers, which can couple CO oxidation with proton reduction and generate H_2_. Physiological characterization has been performed for 32 isolates of (hyper)thermophilic or mesophilic hydrogenogenic CO oxidizers from the five phyla Bacillota, Pseudomonadota, Dictioglomi, Euryarchaeota, and Crenarchaeota (3). Except for *Thermoanaerobacter kivui* and *Carboxydothermus pertinax*, all of these hydrogenogenic CO oxidizers contained *cooS/cdhA* in the gene clusters composed of *cooS/cdhA, cooC*, encoding a maturation factor of CooS (25), *cooF*, encoding a ferredoxin-like protein for electron transfer (26), and genes encoding energy-converting hydrogenases (ECH) (*coo–ech* gene cluster). ECH performs proton reduction, and this process is associated with production of transmembrane ion motive force, which is utilized by the ATP synthase complex for the conversion of ADP to ATP. As a result, Coo/ECH is regarded as the respiratory machinery that supports ATP production (27, 28).

Mo-CODH belongs to the xanthine oxidase family of molybdenum-containing hydroxylases (29, 30). Mo-CODH is divided into two groups: form I Cox and form II Cox. This classification is based on the phylogeny and amino acid sequence motif of the active site of the large subunit CoxL (AYXCSFR for form I, and AYRGAGR for form II). There is also a difference in the organization of *cox* gene clusters in the genome between these two groups (*coxMSL* for form I, and *coxSLM* for form II) (6, 31). Form I CoxL has been well characterized, and its gene has been regarded as a marker of Mo-CODH-mediated CO oxidation (32, 33). CO oxidizers with form I Cox couple CO oxidation with the reduction of quinone (34), and reducing equivalents are finally then to reduce O_2_ in aerobic conditions, or other terminal electron acceptors, such as nitrate and perchlorate, in anaerobic conditions (5, 7, 35, 36). Electron transport generates a proton motive force that contributes to energy conservation (37). Heterotrophic CO oxidizers with form I Cox are assumed to utilize CO, particularly as an energy source, to survive in energy-limited conditions (35). In contrast, some autotrophic CO oxidizers can use CO as both an energy source and a carbon source in the Calvin–Benson–Bassham (CBB) cycle (38, 39). It is uncertain whether all form II Cox-bearing prokaryotes can oxidize CO. Form II Cox purified from Alphaproteobacteria *Bradyrhizobium japonicum* has been demonstrated to oxidize CO (6, 40). CO oxidation has also been observed in a culture of the Bacillota bacterium *Kyrpidia spormannii*, an isolate with form II *cox* but lacking form I *cox* (41). However, isolates from the Alphaproteobacteria family Roseobacteraceae with form II *cox* but no form I *cox* did not exhibit CO oxidation under laboratory conditions (42).

*cooS/cdhA* is commonly found in the genomes of facultative or obligate anaerobes (3), possibly reflecting the sensitivity of Ni-CODHs to O_2_ (43). Form I *coxL* is present in the genomes of aerobes and microaerobes (6, 36). According to our review of the literature, no single prokaryote that can perform both Ni-CODH-mediated and Mo-CODH-mediated CO oxidation has been reported. *cooS/cdhA* and form I *coxL* demonstrate punctate phylogenetic distributions even in various genera. For instance, two of the six species of the genus *Moorella* lack one of the *cooS* homologs, which form the *coo–ech* gene cluster (44). Of the six species of the genus *Aminobacter*, only two possess form I *cox* (45). Such punctate presence and absence patterns can be attributed to horizontal gene transfers or vertical inheritance, followed by recent gene losses (45). The key factors shaping the punctate phylogenetic distributions of *cooS/cdhA* and form I *cox* remain unclear, but could be revealed by a detailed investigation of a monophyletic group, of which some species are CO oxidizers with *cooS/cdhA*, while some are those with form I *cox*.

*Parageobacillus* of the phylum Bacillota is a bacterial genus comprising obligate aerobic or facultatively anaerobic thermophiles (46). Although these bacteria were originally classified as members of the genus *Geobacillus*, they were later reclassified into the recently established genus *Parageobacillus* (47, 48). There are currently six species recognized (*Parageobacillus toebii, Parageobacillus thermoglucosidasius*, *Parageobacillus thermantarcticus*, *Parageobacillus caldoxylosilyticus*, *Parageobacillus galactosidasius*, and *Parageobacillus yumthangensis*), as well as one genomospecies, *Parageobacillus* genomospecies 1 NUB3621 (49). Among them, *P. toebii*, *P. galactosidasius*, and *P. yumthangensis* have recently been proposed as the same species due to high genomic similarity (50). In addition, *Saccharococcus thermophilus*, another species of thermophilic Bacillota bacterium, has been proposed to be re-classified in the genus *Parageobacillus* (50). Isolates of the genus *Parageobacillus* are regarded as ideal candidates to apply in the production of thermotolerant enzymes (51, 52) as well as biofuels and fine chemicals, with higher production efficiency and a lower risk of contamination, because of their thermophilic nature (53–55).

Three *P. thermoglucosidasius* strains, including the type strain (strains DSM 2542^T^, DSM 2543, and DSM 6285) were recently shown to have a *coo–ech* gene cluster and perform hydrogenogenic CO oxidation (56, 57). Furthermore, we isolated another strain of hydrogenogenic CO oxidizer, *P. thermoglucosidasius* TG4, from marine sediments (58). According to subsequent physiological and reverse genetic analyses, the *coo–ech* gene cluster of this species encodes functional Coo/ECH, and both *coo* and *ech* are required for hydrogenogenic CO oxidation (59). Conversely, the type strain of facultatively anaerobic *P. toebii* was demonstrated to have a *coxMSL* gene cluster, corresponding to the typical organization of form I *cox*, though their CO oxidation is yet to be confirmed (56).

In this study, we report that the new isolate G301 from the genus *Parageobacillus* performs both hydrogenogenic and aerobic CO oxidation. We used comparative genomics to expand our knowledge of the physiological diversity, functional versatility, and evolution of CO oxidizers. Additionally, we discussed a factor that contributes to the broad but punctate phylogenetic distribution of CO metabolism.

## Results

### Isolation of strain G301

CO oxidizers were enriched in the sediment of Unagi-ike (31° 13’ 37” N and 130° 36’ 38” E), a freshwater lake in Japan into which water from a nearby hot spring flows, by cultivation at 65 °C under 20% CO and 80% N_2_ atmospheric conditions. Following CO depletion and H_2_ evolution in the gas chromatography, the liquid phase of the enrichment culture was streaked on an NBRC 802 agar plate and incubated at 65 °C in the air. The strain G301 was isolated by selecting one of the colonies that appeared on the plate, indicating that the strain was facultatively anaerobic. G301 cells were rod-shaped, 0.5–1.0 μm×2.0–3.5 μm in size (Fig. 1), and Gram-positive (data not shown). The purity of strain G301 was confirmed by direct sequencing of the partial 16S rRNA gene. The optimal growth temperature and pH were 65 °C and 6.0, respectively, and G301 was tolerant up to 3% NaCl (data not shown). Further cultivation confirmed the hydrogenogenic CO oxidation of G301.

**Fig. 1.**
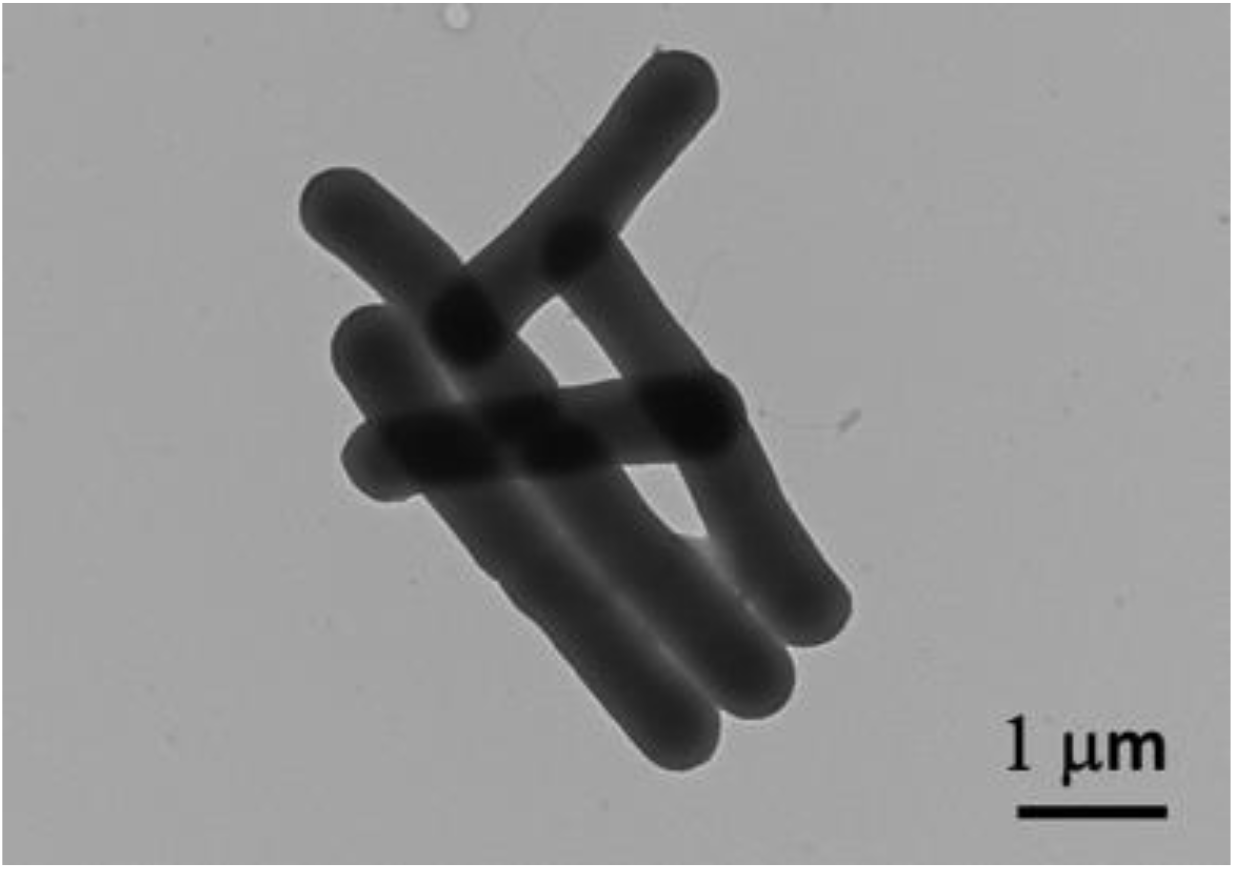
G301 transmission electron microscopy image.

### General features of the G301 genome and phylogenetic relationships with other *Parageobacillus*

The assembled genome of G301 comprised nine contigs with a total length of 3,483,861 bp and a G+C content of 41.79%. Its genome was predicted to contain 13 rRNA genes, 58 tRNA genes, 2,128 protein-coding genes with known functions, and 1,659 functionally unassigned ORFs, according to gene prediction with PATRIC (60).

Although the 16S rRNA sequence of G301 was distantly related to *P. thermoglucosidasius*, the close relatives of G301 did not resolve its phylogenetic position (Fig. 2A). As a result, we performed a phylogenomic analysis with a concatenated dataset of amino acid sequences from 1,505 single-copy core genes of 27 *Parageobacillus* strains, including one strain of *S. thermophilus* and seven strains classified as “unclassified *Geobacillus*” in NCBI taxonomy (61). In this phylogenomic analysis, G301 was sister to the clade of *P. toebii, P. galactosidasius, P. yumthangensis, Geobacillus* sp. 44C, *Geobacillus* sp. NFOSA3, *Geobacillus* sp. E263, *Geobacillus* sp. WCH70 and *Geobacillus* sp. LYN3, with 100% bootstrap support (Fig. 2B). The monophyly containing *P. toebii*, *P. galactosidasius* and *P. yumthangensis* was consistent with a previous study indicating that they were heterotypic synonyms (50). The average nucleotide identity (ANI) values between the genome of G301 and those of *P. toebii, P. galactosidasius, P. yumthangensis*, and the five unclassified *Geobacillus* strains mentioned above ranged from 96.0% to 97.1% (Table 1). These values exceed the 95% threshold for the same species by a significant margin (62). This was consistent with the similarity in general features between the genomes (genome size ranging from 3.22 to 3.80 Mb and G+C content below 43%) (Table 1) comparing to the other *Parageobacillus* strains. Hydrogenogenic CO oxidation has not been reported in any of the strains except G301 whose genomes showed ANI values with G301 ≥95%.

**Fig. 2.**
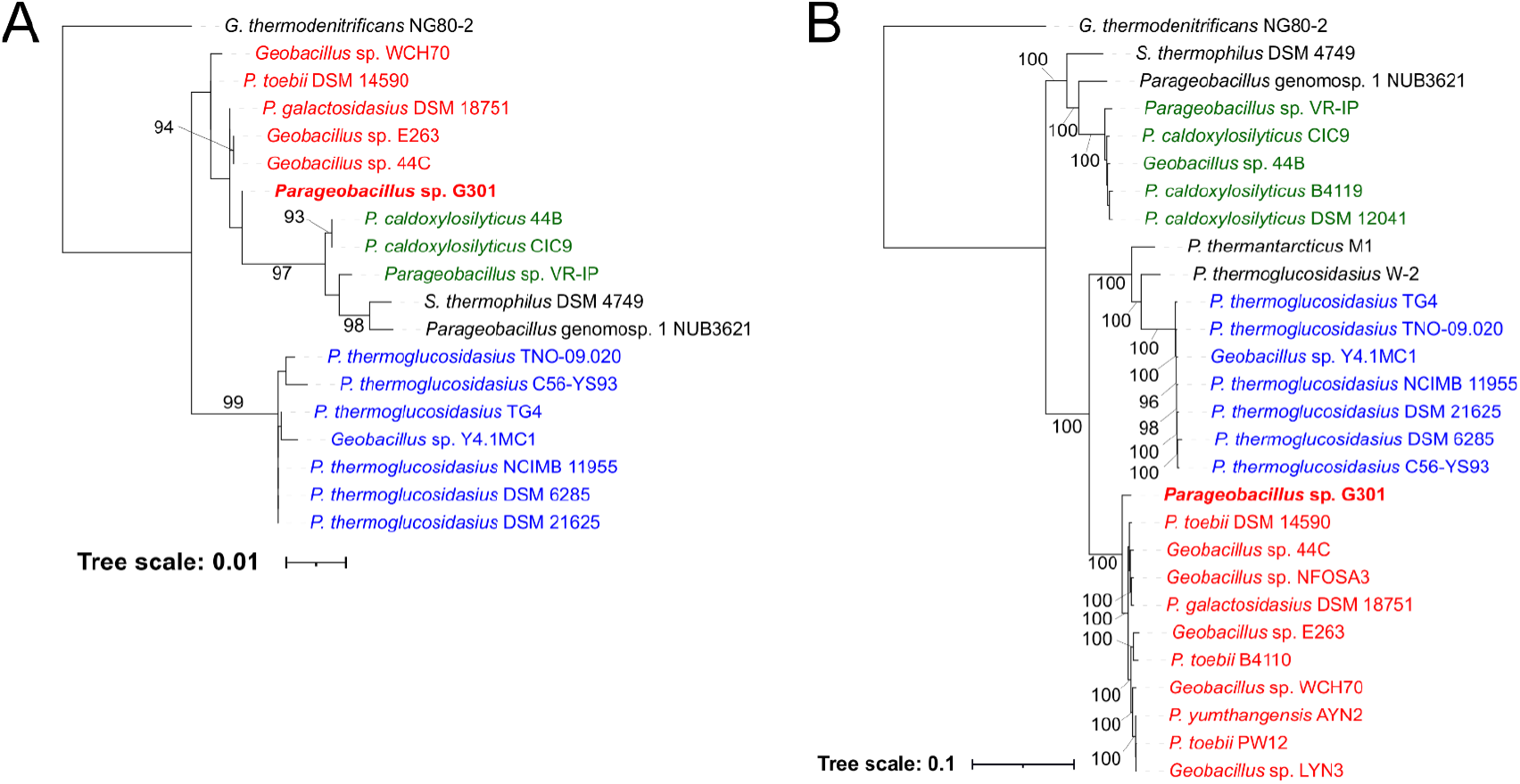
(A) Maximum-likelihood trees of 18 *Parageobacillus* strains and *G. thermodenitrificans* inferred by 16S rRNA gene sequences. (B) maximum-likelihood trees of 27 *Parageobacillus* strains and *G. thermodenitrificans* inferred by the amino acid sequence of 1,505 single-copy core genes. Bootstrap values over 90% are described. Branch lengths represent substitutions per site. Highlighted in red are strains whose ANI values with G301 are higher than 95%. Highlighted in green are strains which show ANI values with *P. caldoxylosilyticus* DSM 12041 higher than 95%. Highlighted in blue are strains which show ANI values with *P. thermoglucosidasius* NCIMB 11955 higher than 95%.

**Table 1.** General features and the ANI values against the genome of G301 of the genomes of 27 *Parageobacillus* isolates

| Strain | Genome size (bp) | G+C (%) | No. of |  | Protein-Encoding<br>Genes with<br>Functional<br>Assignment | Protein-Encoding<br>Genes without<br>Functional<br>Assignment | rRNA | tRNA | Reference | Accession | ANI vs<br>G301 |
| --- | --- | --- | --- | --- | --- | --- | --- | --- | --- | --- | --- |
|  |  |  | Contigs |  |  |  |  |  |  |  |  |
| *Parageobacillus* sp. G301 | 3,483,861 | 41.79 | 9 |  | 2,128 | 1,659 | 13 | 58 | This study | BSDB01000001–<br>BSDB01000009 | 100 |
| *Geobacillus* sp. 44C | 3,220,397 | 42.32 | 121 |  | 2,107 | 1,436 | 9 | 74 | (108) | GCF_002077865.1 | 97.1 |
| *P. toebii* NBRC 107807 | 3,323,060 | 42.36 | 2 |  | 2,127 | 1,476 | 28 | 90 | (109) | GCF_003688615.2 | 97.0 |
| *Geobacillus* sp. NFOSA3 | 3,394,841 | 42.13 | 268 |  | 2,183 | 1,653 | 11 | 91 | (110) | GCF_013096895.1 | 96.8 |
| *P. galactosidasius* DSM 18751 | 3,794,830 | 41.57 | 6 |  | 2,258 | 1,949 | 15 | 95 | (111) | GCF_002217735.1 | 96.7 |
| *P. toebii* PW12 | 3,238,931 | 42.43 | 232 |  | 2,049 | 1,659 | 2 | 61 | (112) | GCF_003367455.1 | 96.3 |
| *P. yumthangensis* AYN2 | 3,409,966 | 42.33 | 124 |  | 2,173 | 1,718 | 4 | 79 | (113) | GCF_002494375.1 | 96.2 |
| *Geobacillus* sp. LYN3 | 3,239,860 | 42.56 | 110 |  | 2,065 | 1,655 | 9 | 83 | (114) | GCF_003064505.1 | 96.2 |
| *P. toebii* B4110 | 3,527,506 | 42.15 | 201 |  | 2,232 | 1,698 | 11 | 91 | (115) | GCF_001587455.1 | 96.2 |
| *Geobacillus* sp. E263 | 3,477,599 | 42.61 | 1 |  | 2,199 | 1,515 | 28 | 92 | (116) | GCF_007197795.1 | 96.1 |
| *Geobacillus* sp. WCH70 | 3,508,804 | 42.80 | 3 |  | 2,179 | 1,536 | 28 | 92 | (117) | GCF_000023385.1 | 96.0 |
| *P. thermoglucosidasius* W-2 | 3,894,562 | 43.15 | 44 |  | 2,399 | 2,037 | 1 | 76 | (118) | GCF_001655645.1 | 85.9 |
| *Parageobacillus* genomsp. 1 NUB3621 | 3,622,385 | 44.38 | 1 |  | 2,250 | 1,681 | 21 | 75 | (119) | GCF_000632515.1 | 85.7 |
| *P. thermantarcticus* M1 | 3,448,881 | 43.67 | 106 |  | 2,192 | 1,758 | 8 | 78 | (120) | GCF_900111865.1 | 85.7 |
| *P. thermoglucosidasius* C56-YS93 | 3,993,793 | 43.93 | 3 |  | 2,446 | 2,070 | 27 | 90 | (121) | GCF_000178395.2 | 85.2 |
| *Geobacillus* sp. Y4.1MC1 | 3,911,947 | 44.01 | 2 |  | 2,424 | 1,971 | 27 | 90 | (122) | GCF_000166075.1 | 85.2 |
| *P. thermoglucosidasius* DSM 6285 | 3,967,726 | 43.58 | 9 |  | 2,377 | 2,121 | 16 | 91 | (57) | GCF_014218645.1 | 85.2 |
| *P. thermoglucosidasius* DSM 21625 | 4,006,039 | 43.83 | 22 |  | 2,444 | 2,109 | 21 | 90 | (123) | GCF_014218665.1 | 85.2 |
| *P. thermoglucosidasius* NCIMB 11955 | 4,004,923 | 43.83 | 3 |  | 2,443 | 2,017 | 27 | 90 | (124) | GCF_001700985.1 | 85.1 |
| *P. thermoglucosidasius* TG4 | 3,946,937 | 43.93 | 11 |  | 2,425 | 2,047 | 25 | 90 | (58) | GCF_003865195.1 | 85.1 |
| *P. thermoglucosidasius* TNO-09.020 | 3,740,238 | 43.86 | 1 |  | 2,465 | 2,342 | 20 | 88 | (125) | GCF_000258735.1 | 85.0 |
| *S. thermophilus* DSM 4749 | 3,145,141 | 44.88 | 3 |  | 1,965 | 1,347 | 26 | 88 | (126) | GCF_011761475.1 | 85.0 |
| *P. caldxylosilyticus* B4119 | 3,949,593 | 44.01 | 144 |  | 2,519 | 1,898 | 7 | 87 | (115) | GCF_001587525.1 | 85.0 |
| *P. caldxylosilyticus* CIC9 | 3,826,547 | 44.17 | 82 |  | 2,462 | 1,679 | 3 | 73 | (127) | GCF_000313345.1 | 84.9 |
| *P. caldxylosilyticus* DSM 12041 | 3,766,511 | 43.96 | 67 |  | 2,414 | 1,699 | 9 | 76 | (126) | GCF_014196025.1 | 84.8 |
| *Geobacillus* sp. 44B | 3,543,185 | 44.62 | 284 |  | 2,354 | 1,601 | 10 | 80 | (108) | GCF_002077755.1 | 84.8 |
| *Parageobacillus* sp. VR-IP | 3,798,106 | 43.96 | 21 |  | 2,433 | 1,784 | 10 | 81 | n/a | GCF_013357975.1 | 84.6 |

### Genes involved in CO oxidation in G301

To characterize the CO oxidation genes in the newly isolated strain G301, we examined key genes of Ni-CODH and Mo-CODH, *cooS* and *coxL*, respectively, in the genome of G301. Notably, our genome investigation revealed that G301 had both these genes. According to the annotation with PATRIC, *cooS* was present in sequence 1 (locus tag: PG301_04030, 373,389–375,311 bp in sequence 1) between *cooC*, encoding a maturation factor of CooS (PG301_04020) (25), and *cooF*, encoding a ferredoxin-like protein involved in electron transfer (PG301_04040) (26) (Fig. 3A). Furthermore, 14 genes encoding the catalytic subunits and maturation factors of hydrogenase (locus tag: PG301_04050–PG301_04160) were in the downstream region of *cooF* (Fig. 3A). According to the HydDB classification (63), the large subunit of hydrogenase encoded by PG301_04100 is a [NiFe] Group 4a hydrogenase. The composition of the *coo–ech* gene cluster in G301 corresponded to that identified in other hydrogenogenic CO oxidizers in a previous study (14, 18) (Fig. 3B).

**Fig. 3.**
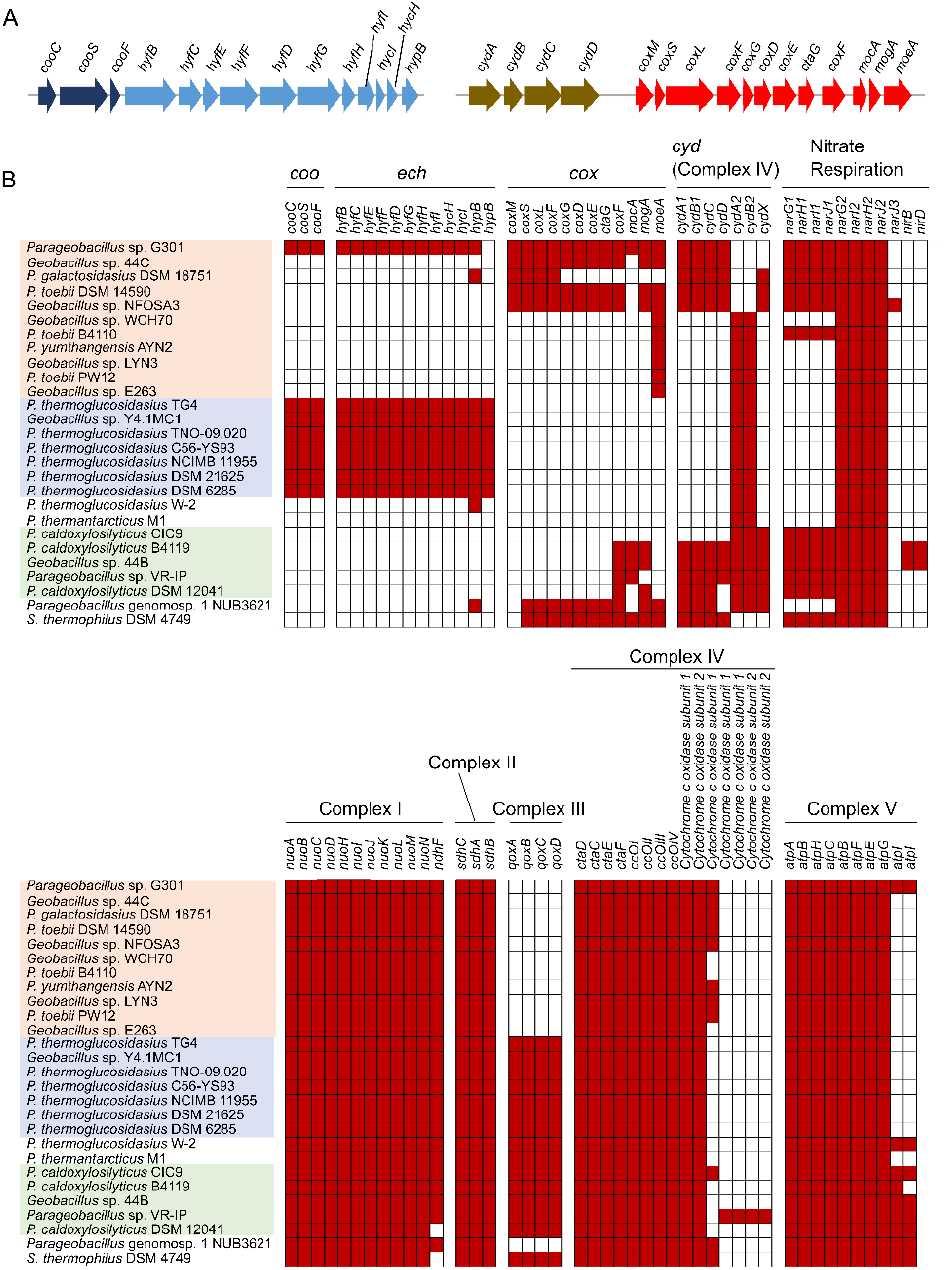
(A) The organization of the *coo–ech* gene cluster and the *cyd–cox* region of G301. Arrows show coding regions and directions. The *coo* gene cluster, *ech* gene cluster, *cox* gene cluster, and *cyd* gene cluster are colored in blue, light blue, dark yellow, and red, respectively. (B) A summary of the presence and absence of *codh* and respiratory genes in *Parageobacillus* genomes that can couple with CO oxidation. Top shows the presence and absence of *coo*, *ech*, *cox*, *cyd*, and *nar* genes, while bottom shows the presence and absence of respiratory complex genes other than *cyd* genes. The names of the strains are colored in the same manner as Fig. 2. Genes filled in red and white are those detected and undetected, respectively. Although *cooF* of *Geobacillus* sp. Y4.1MC1 was not identified in the annotation by PATRIC, it was regarded as present because a *cooF* gene was predicted and annotated in the genome data of RefSeq.

The G301 genome also contained two *coxL* homologs, one of which in the same contig carrying the *coo–ech* gene cluster was identified as form I *coxL* according to the phylogeny and active site motif (locus tag: PG301_05530, 523,179–525,530 bp in sequence 1) (Fig. 3A). *coxM* (locus tag: PG301_05560) and *coxS* (locus tag: PG301_05550), which encode the medium and small subunit of Cox, respectively (15), were also found in the region upstream from *coxL*. Downstream of *coxL*, there were genes encoding CoxF, CoxG, CoxD, CoxE, cytochrome *c* oxidase assembly protein CtaG, CoxF, nucleotidyltransferase family protein MocA, molybdopterin adenylyltransferase MogA, and molybdopterin molybdenum transferase MoeA (locus tags: PG301_05530, PG301_05520, PG301_05510, PG301_05500, PG301_05490, PG301_05480, PG301_05470, PG301_05460, PG301_05450, respectively). The *cox* gene cluster organization, including *coxD, coxE*, and *coxF*, as well as *coxMSL* was reported as a common feature of the form I *cox* gene cluster organization of CO oxidizers with Mo-CODH (6), implying that the genes of G301 are highly likely to encode functional Mo-CODH. Other genes in the form I *cox* gene cluster that encode homologs of accessory proteins of Cox or genes associated with biosynthesis of molybdenum cofactor, may help in the maturation of Cox catalytic subunits. Additionally, another gene cluster comprising genes that encode cytochrome ubiquinol oxidase subunits CydA and CydB (locus tags: PG301_05600 and PG301_05590) and their maturation factors CydC and CydD (locus tags: PG301_05580 and PG301_05570) was present upstream of *coxM* (Fig. 3A). The *cyd* gene cluster was separated from the *cox* gene cluster by a 1,657 bp-long non-coding region. The gene content and order of *cyd* were similar to those reported for the *cyd* operon in *Bacillus subtilis* (64). The arrangement of *cyd* and *cox* gene clusters (hereafter stated *“cyd–cox* region”) suggests that they are functionally related.

### Respiratory machineries that can couple with CO oxidation

To estimate the physiological functions of CO oxidation in G301, we conducted a comprehensive investigation of the draft genome of G301 for the respiratory machineries, some of which might receive electrons from CO oxidation. G301 had genes encoding essential parts of the aerobic respiratory chain, namely *nuoABCDHIJKLM* encoding Complex I, *sdhABC* encoding Complex II, as well as genes encoding terminal oxidases such as CydABCD, CtaDCEF, cytochrome *c* oxidase subunits I and II, and CcoI–IV (Fig. 3B). Regarding anaerobic respiratory machineries, two gene clusters of the respiratory nitrate reductase (*narG1H1I1J1* and *narG2H2I2J2*) were found (Fig. 3B). Both aerobic respiration and respiratory nitrate reduction utilize quinones for electron transfer (65, 66). CO-derived electrons by Mo-CODH may be received by quinones (34), implying that nitrate reduction, in addition to aerobic respiration, is a candidate respiratory machinery coupled with Mo-CODH-mediated CO oxidation by G301. Quinone-dependent respiratory machinery, except the aerobic respiratory chain or nitrate reductase, was not identified in this study. The gene for ribulose 1,5- bisphosphate carboxylase/oxygenase that is responsible for CO_2_ fixation in the CBB cycle was not detected in the G301 genome, indicating that Mo-CODH-mediated CO oxidation is unlikely to be coupled with CO_2_ fixation, as reported in other prokaryotes (67). For Ni-CODH, *rnf* responsible for the electron transfer from ferredoxin to NAD^+^ (68) was not identified in the G301 genome. Electrons obtained from CO by Ni-CODH re transferred to CooF or ferredoxin, and used for further oxidation-reduction reactions, such as H_2_ evolution by ECH (69) and NAD^+^ reduction to NADH by Rnf (70), which can be used for anaerobic respiration (3). It appears that G301 use ECH to accept electrons from Ni-CODH-mediated CO oxidation.

### Distribution of *coo–ech* gene clusters and *cyd–cox* regions in *Parageobacillus* genomes

To explore the distribution of *coo–ech* gene clusters and *cyd–cox* regions in the genus *Parageobacillus*, we comprehensively examined those genes in available *Parageobacillus* genomes by Orthofinder and PATRIC.

A single copy of the *cooS* was present in the genomes of all the analyzed genomes which showed ANI value ≥95% with that of *P. thermoglucosidasius* NCIMB 11955 (Fig. 2B). To the best of our knowledge, *coo–ech* of NCIMB 11955 and TNO-09.020 had never been characterized before. The genes of *coo–ech* were conserved among all of the analyzed genomes with *cooS* (Fig. 3B). In contrast, the form I *cox* was found in various species/strains of the genus *Parageobacillus* rather than *coo–ech*, including *Parageobacillus* genomosp. 1 NUB3621, *S. thermophilus* DSM 4749, *P. toebii* DSM 14590, *Geobacillus* sp. 44C, *Geobacillus* sp. NFOSA3, and *P. galactosidasius* DSM 18751. However, it is unclear whether the Cox of *P. galactosidasius* DSM 18751 is as functional as those of strains with the entire set of maturation factor genes because it lacks many genes encoding putative Cox maturation factors. The *cyd–cox* regions were also present in all the genomes having form I *cox*, as seen in G301 (Fig. 3A). No publicly available *Parageobacillus* genome have both *cooS* and form I *cox*, indicating that G301 is unique in the genus *Parageobacillus*.

To determine whether there was another prokaryotic isolate that can use CO both aerobically and anaerobically, as predicted for G301, we investigated both *cooS/cdhA* and form I *coxL* against all the prokaryotic isolate genomes (183,204 genomes) in the RefSeq/Genbank genome database. As a result, while 4,305 and 10,540 isolates had *cooS/cdhA* and *coxL*, respectively, only six had both *cooS/cdhA* and *coxL* (Fig. 4A). Moreover, none of the six genomes had both *cooS/cdhA* and form I *coxL*, further highlighting the uniqueness of G301 (Fig. 4B).

**Fig. 4.**
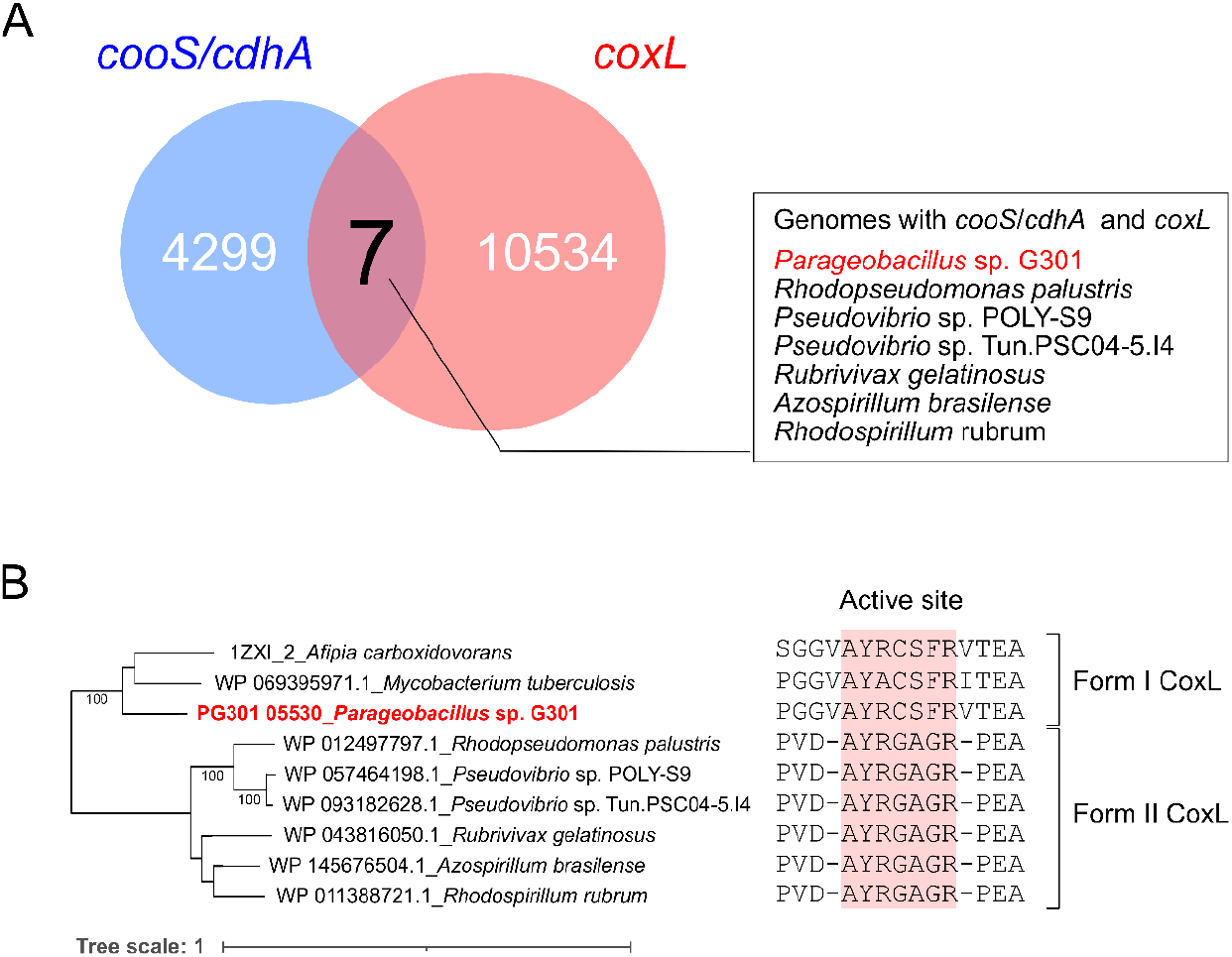
Distributions of *cooS/cdhA* and *coxL* in isolate genomes. (A) Benn diagram of the numbers of isolates with *cooS/cdhA* and isolates with *coxL*. blue and red circles represent the number of genomes with *cooS/cdhA* and those with *coxL*, respectively. In addition to G301, 183,204 isolate genomes in GenBank were analyzed. (B) Phylogenetic tree and the active site motif of CoxL of isolates containing both *cooS/cdhA* and *coxL*. Bootstrap values >90% are described. The tree scale represents substitutions per site. CoxL of *M. tuberculosis* and *A. carboxidovorans* is known as Form I CoxL.

### Genomic traits exclusively shared among *Parageobacillus* strains having the *coo–ech* gene cluster or *cyd–cox* region

The respiratory machineries found in the G301 genome is broadly distributed among *Parageobacillus* genomes. Additionally, comparative genomic analysis revealed that all the examined strains had the entire set of aerobic respiratory chains (Fig. 3B).

However, there were two distinct sets of *cydA* and *cydB*, referred to as *cydA1B1* and *cydA2B2*, the former of which was included in the *cyd–cox* region (Fig. 3B). The *cydA1B1* gene was also identified in the four of five genomes which showed ANI value ≥95% with *P. caldoxylosilyticus* DSM 12041 that did not have the form I *cox* gene cluster. Except for *Parageobacillus* genomosp. 1 NUB3621 and *P. caldoxylosilyticus* DSM 12041 lacking *cydD*, all genomes with *cydA1B1* also had *cydCD*. The *cydA2B2* was present in the genomes of *Parageobacillus* genomosp. 1 NUB 3621, and all the form I *coxL*-lacking strains (Fig. 3B). In addition, there were two distinct gene clusters for nitrate reductase, *narG1H1I1J1* and *narG2H2I2J2*, as stated above. The *narG1H1I1J1* gene was found in 12 out of 27 genomes, but it was present in all form I *cox*-containing strains. The *narG2H2I2J2* gene was found in all of the *Parageobacillus* strains analyzed (Fig. 3B). The absence of *rnf was* also observed in all of the genomes examined.

We conducted comparative genome analyses for *Parageobacillus* genomes to explore if *codh* retention could affect genomic structures and if certain cellular functions unrelated to respiration could also couple with CO oxidation. We first compared whole-genome synteny among *Parageobacillus* sp. G301, *P. toebii* DSM 14590, and *Geobacillus* sp. WCH70, which are all closely related but have distinct sets of *codhs* (Figs. 2A, 2B, and 3B). No explicit genomic differences were detected among the three strains on this scale. The genome synteny of these strains was highly conserved, indicating that the presence or absence of *coo–ech* gene clusters and *cyd–cox* regions would not result in any unique whole-genome rearrangements (Fig. 5A, 5B, and 5C). Subsequently, we compared the synteny of genomic regions around the *coo–ech* gene clusters and *cyd–cox* regions. Conserved genes were present in and around *coo–ech* gene clusters of the analyzed genomes (Fig. 5D). Conservation of synteny around the *cyd–cox* regions was also observed, although with minor differences in the form I *cox* and *cyd* gene clusters, such as translocation or addition of genes (Fig. 5E). The genes surrounding the gene clusters were also close to each other in *codh-lacking* genomes. The *coo–ech* gene clusters, as well as *cyd–cox* regions, were found in homologous loci in the *Parageobacillus* genomes. Ultimately, we investigated the genes uniquely encoded by genomes with *coo–ech* gene clusters and those with *cyd–cox* regions. As a result, only one gene, except for genes encoded in the *coo–ech* gene cluster, was uniquely present in the genomes with *coo–ech* (Fig. 6). This gene encodes undecaprenyl-diphosphatase UppP, which contributes to the biosynthesis of the cell wall (71). However, it remains unclear whether the encoded protein is involved in or affected by hydrogenogenic CO oxidation. Conversely, no genes, except for genes in the *cox* gene cluster and the *cyd* gene cluster, were uniquely identified in genomes with *cyd–cox* regions (Fig. 6).

**Fig. 5.**
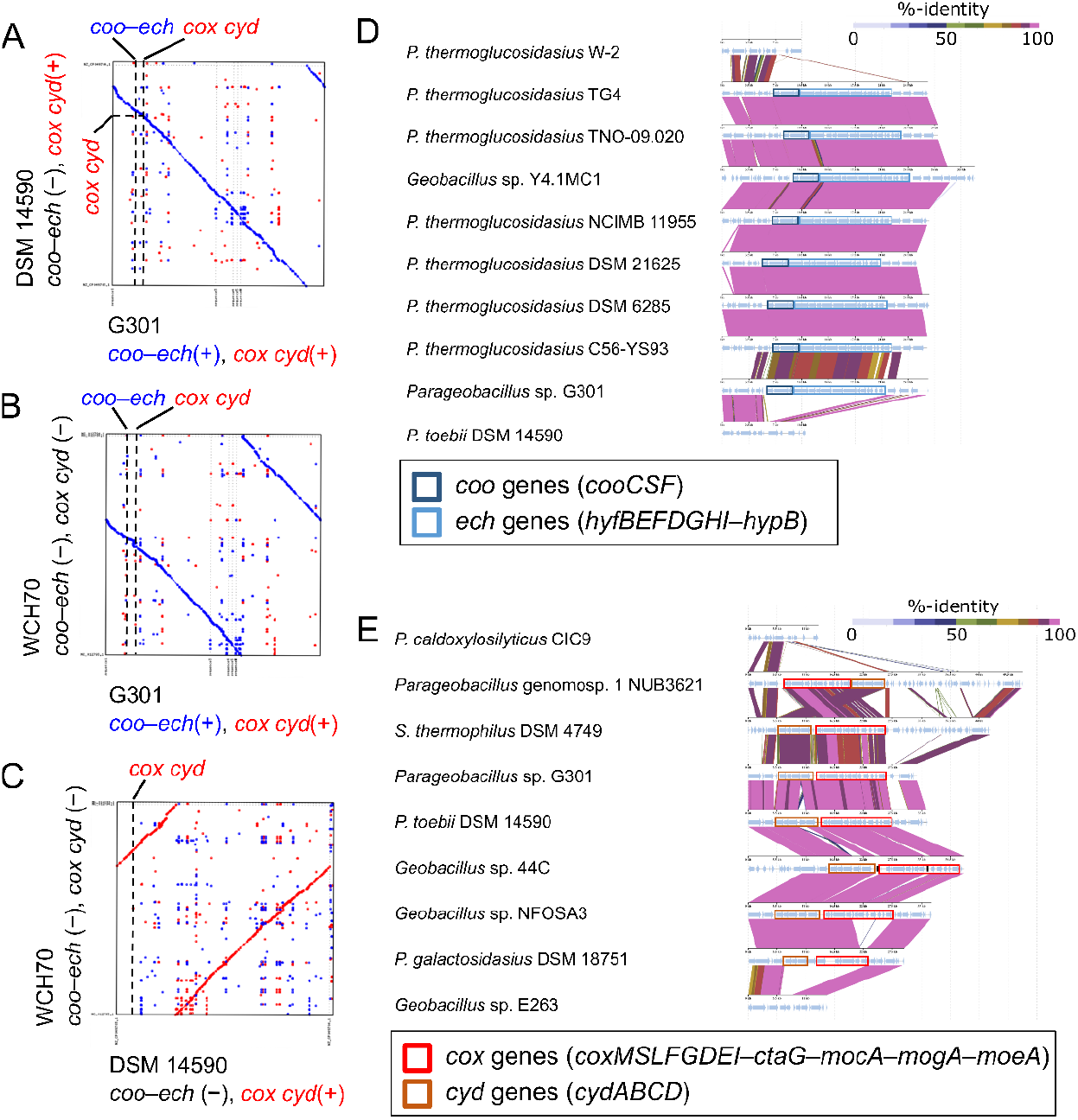
Synteny comparisons of *Parageobacillus* genomes with and without *codhs*. Whole genomes of (A) *Parageobacillus* sp. G301 and *P. toebii* DSM 14590, (B) *Parageobacillus* sp. G301 and *Geobacillus* sp. WCH70, and (C) *P. toebii* DSM 14590 and *Geobacillus* sp. WCH70. For (A)–(C), plus and minus symbols in both axes indicate the presence and absence of *coo–ech* and *cyd–cox*, respectively. (D) Gene synteny of *coo–ech* gene clusters and their flanking regions in the *Parageobacillus* genomes. Seven genomes of *P. thermoglucosidasius* and one each of *P. toebii* and *Geobacillus* sp. were compared in addition to G301. (E) Gene synteny of *cyd–cox* regions in the *Parageobacillus*. the black lines on the *cyd–cox* region of *Geobacillus* sp. 44C indicate the division of the corresponding genomic region into separate contigs.

**Fig. 6.**
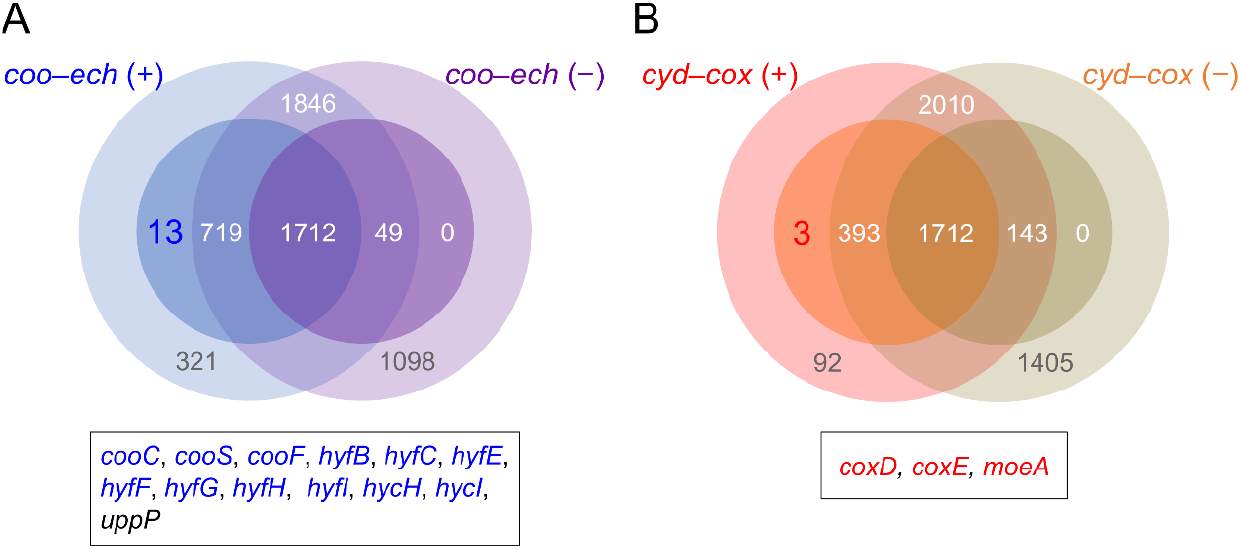
Comparison of the core and pan genomes of *Parageobacillus* genomes with and without *codh* gene clusters. (A) Core and pan genomes of *Parageobacillus* genomes with *coo–ech* gene clusters (*coo–ech* (+)) and without *coo–ech* gene clusters (*coo–ech* (–)). The inner circle represents the core genomes, while the outer circle represents the pan genomes. The number indicates shared genes in each category. The category for genes shared exclusively by all the *coo–ech* gene cluster-containing genomes is highlighted, and those genes shown in the box are colored blue if they are located in the *coo–ech* gene clusters. (B) Core and pan genomes of *Parageobacillus* genomes with *cyd–cox* regions including accessory genes of *cox (cox, cyd* (+)), and without either or neither of *cox* and *cyd* gene clusters (*cox, cyd* (–)). the category for genes shared exclusively by all the genomes with *cyd–cox* region is highlighted, and those genes shown in the box are colored red if they are located in the *cox* and *cyd* gene clusters. Other details are described in (A).

Overall, the retention of CO oxidation genes in *Parageobacillus* genomes would have a minor effect on genomic structures and cellular functions other than CO-mediated electron transport.

### CO metabolisms of G301 cultures

We characterized the CO metabolism of G301 by cultivating it in the presence or absence of CO with different terminal electron acceptors, *i.e*., proton, O_2_, and nitrate, and with yeast extract as a carbon source. G301 was confirmed to consume CO in all the tested conditions (Fig. 7A, 7D, and 7G). In the cultures with CO but without O_2_ and nitrate, CO consumption as well as H_2_ and CO_2_ production were observed from the late log phase to the stationary phase (4–24 h after inoculation) (molar ratio of consumed CO: H_2_ evolved in the gas phase: CO_2_ evolved in the gas phase = 1:1.03:0.47) (Fig. 7A). No H_2_ or CO_2_ production was observed in the same condition, except for the absence of CO (Fig. 7B). Growth yield with CO was significantly higher than that without CO (*p* = 0.025) (Fig. 7C). CO most likely supported growth in this condition, possibly as an energy source, as indicated in previous studies (27). This result was in line with the phenomena observed for *P. thermoglucosidasius* (59).

**Fig. 7.**
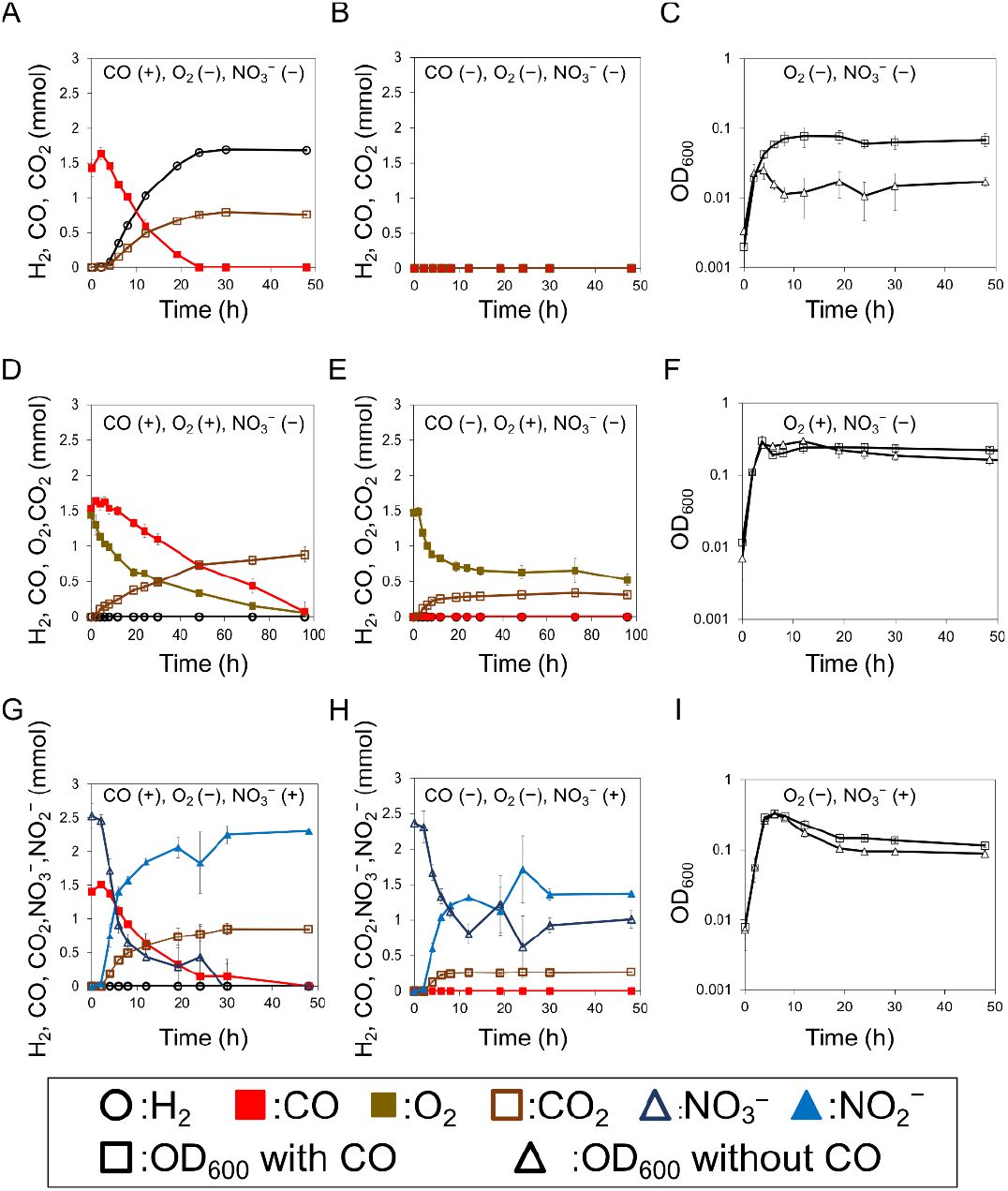
Growth of the G301 cultures, and the levels of CO, CO_2_, H_2_, O_2_, nitrate, and nitrite. The presence and absence of CO, O_2_, and nitrate in the initial conditions are shown in parenthesis in each panel. Plus and minus symbols indicate presence and absence, respectively. (A) Amount of H_2_, CO, and CO_2_ in the culture with CO but without O_2_ and nitrate. (B) amount of H_2_, CO, and CO_2_ in the culture without CO, O_2_, and nitrate. (C) growth of cultures under conditions A and B. (D) amount of O_2_, CO, and CO_2_ in the culture with CO and O_2_ but without nitrate. (E) amount of O_2_, CO, and CO_2_ in the culture with O_2_ but without CO and nitrate. (F) growth of cultures under the conditions of D and E. (G) amount of CO, CO_2_, nitrate, and nitrite in the culture with CO and nitrate but without O_2_. (H) amount of CO, CO_2_, nitrate, and nitrite in the culture with nitrate but without CO and O_2_. (I) growth of cultures under the conditions of H and I. plots represent the means of three biological replicates. Thin vertical lines represent standard deviations.

In the presence of CO and O_2_, the consumption of CO and O_2_ and the production of CO_2_ were observed until 96 h (molar ratio of consumed CO: consumed O_2_: CO_2_ evolved in the gas phase= 1:0.95:0.60), while no H_2_ production was observed (Fig. 7D). The levels of consumed O_2_ and CO_2_ evolved in the gas phase at 96h after inoculation were 32% and 64% lower in the culture without CO, respectively than in the culture with CO, implying that G301 performed aerobic CO oxidation (Fig. 7E). Growth rate in the log phase (0–4 h) as well as growth yield in the culture with CO were not significantly different from those without CO (*p* = 0.19 for growth rate, and *p* = 0.33 for growth yield) (Fig. 7F). However, the depletion of OD_600_ in the stationary phase tended to be slower in the culture with CO (Fig. 7F).

In anaerobic cultures with CO and nitrate, complete consumption of CO and nitrate, as well as production of CO_2_ and nitrite, was observed after 48 h (molar ratio of consumed CO: consumed nitrate: CO_2_ evolved in the gas phase: produced nitrite = 1:1.79:0.60:1.63). No H_2_ production was observed as in the presence of O_2_ (Fig. 7G). The amounts of consumed nitrate, CO_2_ evolved in the gas phase, and produced nitrite at 96h after inoculation were 46%, 69%, and 40% lower in the absence of CO than in the presence of CO (Fig. 7H), indicating the presence of CO oxidation coupled with nitrate reduction. Growth rate in the log phase (0–4 h) and growth yield in CO-treated cultures were not significantly different from those in non-CO-treated cultures (*p* = 0.90, growth rate and *p* = 0.66 for growth yield). The depletion of OD_600_ in the stationary phase tended to be slower in the presence of CO, as observed in aerobic CO oxidation (Fig. 7I).

To determine which of the CODHs in G301 was responsible for CO oxidation in the presence of nitrate, we examined the CO oxidation of *P. thermoglucosidasius*, which had a CooS with over 70% amino acid identity to that of G301 but lacked *cox*. *P. thermoglucosidasius* demonstrated CO consumption, H_2_ production, and CO_2_ production in cultures with KCl instead of nitrate as the control (Fig. S1A). However, hydrogenogenic CO oxidation was not observed in nitrate-containing cultures (Fig. S1B), implying that *P. thermoglucosidasius* did not couple nitrate reduction and Ni-CODH-mediated CO oxidation at detectable levels.

## Discussion

Sequencing, *de novo* assembly, and comparative genomic analyses for *Parageobacillus* sp. G301 revealed that this strain is the only isolate that has both *cooS/cdhA* and form I *cox* (Fig. 4). Cultivation experiments validated the genome-based estimation for CO oxidation in G301, both aerobically and anaerobically (Fig. 7), indicating that both Ni-CODH and Mo-CODH were functional in this strain. In particular, no Ni-CODH-mediated CO oxidation coupled with nitrate reduction was identified in *P. thermoglucosidasius*, indicating that the CO oxidation coupled with the nitrate reduction in G301 was most likely mediated by Mo-CODH rather than Ni-CODH, whose amino acid sequence was highly similar to that of *P. thermoglucosidasius*. Overall, G301 is the sole isolate that is capable of performing CO oxidation coupled with H_2_ production and O_2_ reduction, as well as nitrate reduction. Thus, G301 would gain energy with CO-mediated respiration in disparate environmental conditions. This metabolic trait may be advantageous in an oxic-anoxic interface because it allows prokaryotes to use CO from the atmosphere or other sources such as hydrothermal vents (72), soil (73), seawater (74) or freshwater environments (75, 76) with fluctuating O_2_ levels. Explorations in such changeable environmental conditions would lead to the discovery of novel CO oxidizers with previously unknown combinations of distinct CO metabolisms, providing further insight into the diversity, evolution, and industrial applications of such prokaryotes with intriguing respiration.

G301 further allowed us to investigate a factor that contributed to the broad but punctate phylogenetic distribution of CO metabolism. Comparative genomic analyses revealed that each of hydrogenogenic CO oxidation with Ni-CODH, anaerobic CO oxidation coupled with nitrate reduction with Mo-CODH, and aerobic CO oxidation with Mo-CODH showed punctate but not ubiquitous phylogenetic distribution in the genus *Parageobacillus*. The punctate distribution of CO metabolisms might be explained by the lack of significant influence of *codh* gene clusters on cellular functions, given that there are no obvious explicit differences in genomes and gene repertoires between CO oxidizers and non-CO oxidizers (Fig. 5). This is consistent with our previous observation that *P. thermoglucosidasius* mutants lacking *codh* could grow with or without CO (59). CODHs genes might act as an optional genetic element that improve fitness in limited conditions rather than essential genes for cell viability. If this is the case, gains and losses of CODH genes may have had less of an impact on cell physiology and viability; as a result, the current punctate phylogenetic distribution of *codh* genes in this genus may have been shaped. Other than *Parageobacillus*, there are taxa with a punctate distribution of *codh* genes, and CO oxidation does not appear to make a significant difference in overall cellular functions. In the family Roseobacteraceae, a group of marine bacteria formally known as the Marine Roseobacter Clade (MRC), CO oxidizers with form I *cox* gene cluster and non-CO oxidizers without form I *cox* gene cluster are closely related (42). An aerobic CO oxidizer of this family, *Ruegeria pomeroyi*, exhibited no observable difference in growth or metabolome between cells grown with and without CO (77). Another example is *Thermoanaerobacter*, a genus of thermophilic anaerobic acetogens. Two anaerobic hydrogenogenic CO oxidizer strains and one strain with the *coo– ech* gene cluster have been characterized, while no other strains with *coo–ech* are known (11, 44, 78). *T. kivui*, a hydrogenogenic CO oxidizer, had been deleted one of the two *cooS* genes which was responsible for hydrogenogenic CO oxidation. The resulting mutant lacked the ability to grow on CO, but it grew similarly to the parental strain on glucose, mannitol, H_2_+CO_2_ or formate (79). If their *codh* genes are also optional genetic elements, the punctate distribution of *codh* and CO metabolisms in various prokaryote lineages may also be explained, at least in part, as for *Parageobacillus*.

## Materials and methods

### Sampling

Sediment samples were collected from Unagi-ike (31° 13’ 37” N, 130° 36’ 3’’ E) on May 12, 2,018. The collected sample was preserved in a 50 mL plastic tube, and the headspace was filled with N_2_ gas (GL Science Inc.). The sediment was kept anaerobic by packing it in a freezing bag with Anaero Pack (Mitsubishi Gas Chemical), cooling it with ice during transportation to the laboratory, and storing it at 4 °C until use.

### Isolation

As liquid medium for enrichment culture, we used the modified B medium (59) from the B medium used in the previous study (9); the medium was comprised of 0.03 g Na_2_SiO_3_, 0.5 g NH_4_Cl, 0.1 g KH_2_PO_4_, 0.2 g MgCl_2_·6H_2_O, 0.1 g CaCl_2_·2H_2_O, 0.3 g KCl, 0.1 g NaHCO_3_, 1 g yeast extract, 0.5 mL trace mineral solution SL-6 (80), and 1 mL vitamin solution (81) per litter. For plate cultivation, NBRC 802 agar (82) was used; the medium was composed of 10 g/L hipolypepton, 2 g/L yeast extract, 1 g/L MgSO_4_·7H_2_O, and 15 g/L agar. First, 1 g of the sediment was transferred into the modified B medium and incubated under a 20% CO/80% N_2_ atmosphere at 65 °C in a forced convection oven DRS620DA (Advantec). The gas composition of the enrichment culture was measured by sampling 1 mL of the gas phase into a glass vial SVG-3 (NICHIDEN RIKA-GLASS) filled with air and sealed with a butyl rubber and melamine cap. Subsequently, 0.5 mL of the gas in the vials was analyzed using a GC-2014 gas chromatography system (Shimadzu) equipped with a thermal conductivity detector and a Shincarbon ST-packed column (Shinwa Chemical Industries), using N_2_ as the carrier gas. When consumption of CO and production of H_2_ were observed, the liquid phase of the culture was spread over the NBRC 802 agar plate and incubated at 65 °C overnight under the aerobic condition. One of single-colonies was picked and named strain G301. The purity of the isolate was confirmed by the Sanger sequencing of the PCR-amplified, partial 16S rRNA gene as described in previous literature (83). A determined partial sequence of 16S rRNA was used as the query for a BLASTn search against the NRBI RefSeq genome database to search for close relatives of G301.

### Observation of cell morphology

Aerobically grown G301 cells were harvested during the log phase. After fixation with 25% glutaraldehyde, the cells were negatively stained with 2 % uranyl acetate (84) and observed using transmission electron microscope (H-7650, Hitachi).

### Characterization of growth

Optimal growth conditions were analyzed in aerobic culture using NBRC 802 medium (85) as the liquid phase. Growth was monitored by measuring the OD_600_ value of the culture using an Ultrospec2100 (Amersham) every 30 min, and the condition with the shortest doubling time was regarded as optimal. To determine optimal temperature, cells were G301 cells were cultured at 65 °C and transferred to the fresh medium set at 40, 50, 60, 65, and 70 °C. To determine optimal pH, temperature was set at 65 °C and the pH of the medium was varied in 0.5 increments from 5.5 to 8.5. As buffers for pH adjustment in media with NaOH or HCl, 500 mM MES (for cultures of pH 5.5, 6.0, 6.5, and 7.0), HEPES (for cultures of pH 7.0, 7.5, and 8.5), and TAPS (for cultures of pH 8.0) were used. Tolerance to NaCl was tested in the presence of 0, 3, and 5% NaCl at 65 °C and pH 6.0. Before the assessments of optimal pH and tolerance to NaCl, Precultures were conducted in the same conditions to that of main cultures.

### Genome sequencing

DNA was extracted from G301 using a DNeasy blood and tissue kit (Qiagen). The sequence library was prepared with Nextera Mate Pair Sample Prep Kit (Illumina) according to the manufacture’s protocol, and then sequenced with the MiSeq platform using a MiSeq Reagent Kit v3 (2×300 bp paired-end) (Illumina), yielding 3,272,078 paired-end reads. Quality trimming and adapter removal were performed with Trimmomatic v. 0.3.6 (86) by setting “lluminaclip option” as 2:30:10, “leading” and “Trailing” options as 3, “Slidingwindow” option as 4:15, and “MINLEN” option as 30. Further trimming of mate-pair adapter sequences was performed by NxTrim v. 0.4.1 (87) with the default parameters, resulting in 2,769,352 paired-end reads. The qualified paired-end reads were assembled using SPAdes v. 3.13.0 using the default parameters (88). Mate pair sequences were mapped against assembled contigs using Burrows-Wheeler Aligner (BWA) v. 0.7.17 (89) and SAMtools v. 0.1.19 (90) with the default parameters, and the quality of assembled scaffolds was evaluated by NxRepair v. 0.13 (91). The obtained draft genome was annotated with DFAST v. 1.2.6 (92).

### Phylogenetic analysis of *Parageobacillus* isolates

The genome sequence of G301 and *Parageobacillus* strains retrieved from NCBI RefSeq (93), GTDB (94), and PATRIC (60) were subjected to an all-versus-all comparison of ANI using FastANI (62) with the default parameters. A genome pair that showed an ANI above 99.9% was regarded as identical, and one of the genomes was removed at this step. The genome size, GC content, and number of contigs of remaining genomes were evaluated with quast v. 5.0.2 (95). The number of protein-coding genes was estimated using PATRIC (60).

To determine the phylogenetic position of G301, 16S rRNA gene sequences were retrieved from 18 *Parageobacillus* genomes as well as from the genome of *Geobacillus thermodenitrificans* NG80-2 as an outgroup. Sequences were aligned with the L-INS-I method of MAFFT (96) v. 7.471 and trimmed using trimAl v. 1.4.1 (97) with the automated1 option, resulting in a dataset of 19 taxa and 1,562 sites. The maximum likelihood tree was reconstructed using IQTREE v.1.6.12 (98) under the TN+F+R2 model, which was selected as the best model based on BIC for the dataset, with 1,000 ultrafast bootstrap analyses. The tree was visualized using the Interactive Tree Of Life (iTOL) web server (99). We determined the phylogenetic relationship of *Parageobacillus* strains. based on core protein-coding genes, which were single-copy genes present in all 27 *Parageobacillus* and *G. thermodenitrificans* genomes. All encoded proteins, including hypothetical proteins, were clustered using Orthofinder v. 2.5.2 (100). The translated amino acid sequences of the identified 1,505 core genes in the 28 genomes were aligned with the muscle v.3.8.31 (101) with the default parameters, trimmed with trimAl v. 1.4.1 (97), and concatenated, yielding a dataset of 28 taxa and 421,717 sites. The maximum likelihood tree was reconstructed with IQTREE v. 1.6.12 (98) under the JTT+F+R4 substitution model and selected as the best model based on BIC for the dataset, with 100 bootstrap analyses. The tree was visualized as described above.

### Identification of genomes with *codh* from *Parageobacillus* isolates and from the NCBI genome database

Using CooS of *P. thermoglucosidasius* strain TG4 as a cue, homologs of *cooS* in the analyzed genomes were identified using Orthofinder v. 2.5.2 (100). The obtained protein sequences were confirmed as CooS by phylogenetic analysis, followed by a comparison of the active-site motifs. We constructed a phylogenetic tree with amino acid sequences of CooS homologs with representative CooS sequences of each clade of A to G, as well as mini-CooS (18) (WP_011305243.1, WP_026514536.1, WP_039226206.1, WP_011342982.1, WP_012571978.1, WP_011343033.1, OGP75751.1, and WP_007288589.1, respectively) mentioned in the previous study (14), and hydroxylamine reductase (WP_010939296.1) as the outgroup. These sequences were aligned, and the alignment was trimmed as described above for the phylogenetic analysis of 16S rRNA, resulting in a dataset of 17 taxa and 613 sites. The dataset was subjected to maximum likelihood analysis with IQTREE v. 1.6.12 (98) under the Blosum62+G4 model, which was automatically selected as the best model based on BIC for the dataset, with 1,000 ultrafast bootstrap analyses. The tree was visualized as described above. For identification of the active site motif of CooS homologs, alignment was visualized with MEGA-X (102) and the presence of the conserved catalytic site and surrounding metal clusters described before (14) were checked manually.

Homologs of CoxL were identified as described for CooS, by using CoxL of *P. toebii* strain DSM 14590 as a query. Different from CooS, homologs identified as CoxL might be comprised of different proteins of the xanthine oxidase family (29). We performed phylogenetic analysis followed by comparison of the active site motifs to remove proteins other than form I CoxL. We compared all the detected CoxL homologs with the form I CoxL sequences of *Afipia carboxidovorans* and *Mycobacterium tuberculosis* (1ZXI_2 and SRX92445.1, respectively), both of which were known to oxidize CO by cultivation experiments (103, 104). An enzymatic analysis for *A. carboxidovorans*, in particular, confirmed CO oxidation by the form I CoxL (34). The sequences were aligned and trimmed as described above, resulting in a dataset of 15 taxa and 819 sites. The dataset was subjected to maximum likelihood analysis with IQTREE v. 1.6.12 (98) under the LG+I+G4 model, which was automatically selected as the best model based on BIC for the dataset, with 1,000 ultrafast bootstrap analyses. The tree was visualized as previously described. Alignment was visualized using MEGA-X (98), and the motif of the active site, AYXCSFR, of functional form I CoxL reported in a previous study (6) was checked manually. We removed six proteins that were found in *Parageobacillus* genomes but did not conserve the active site motif and formed a different group from the known form I CoxL on the phylogenetic tree (Fig. S2).

Genomes having both *cooS/cdhA* and form I *coxL* were also explored for available genomes as described in our previous study (19). Briefly, CooS/CdhA was searched using the BLASTp search of DIAMOND v. 0.9.29 (105) against the NCBI non-redundant protein database (February 2,020) using representative sequences from clades A to G of CooS/CdhA and mini-CooS as queries. Sequences with an e-value <0.001, ≥400 amino acids, and including the conserved catalytic site and the surrounding metal clusters described previously (14) were regarded as genuine CooS/CdhA. To identify genomes with form I *coxL*, we searched for genomes carrying *coxL* homologs by BLASTp search using DIAMOND v. 0.9.29, using the amino acid sequence of form I CoxL of *A. carboxidovorans* (1ZXI_2) as a query. Sequences with e-values <0.001, ≥400 amino acids, and bitscores ≥500 were obtained, regardless of whether they had the active site motif of form I CoxL. Genomes containing *cooS/cdhA* and *coxL* were obtained from the RefSeq/GenBank genome database, as described by Omae et al. (2019). CoxL homologs encoded in the genomes of CooS homologs were classified into form I CoxL and other proteins, as described above.

### Comparative genomic analysis

In order to analyze the genomic context of *cooS* and form I *coxL* and to explore the conservation of respiratory machineries, protein coding genes in the *Parageobacillus* genomes were predicted and re-annotated with PATRIC v. 3.6.8 and BLASTp search against NCBI nr protein database. Whole genome synteny was evaluated using the nucmer method of MUMmer v. 3.23 (106). Conservation of genomic region around Ni-CODH/Mo-CODH was compared using ViPTree v. 2.0 (107). The conservation surrounding *coo–ech* was examined by comparing genomes bearing *coo–ech* with each other and with strains without *coo–ech (P. toebii* DSM 14590 for *Parageobacillus* sp. G301, and *P. thermoglucosidasius* W-2 for *P. thermoglucosidasius* TG4). The conservation around *cyd–cox* region was analyzed by comparing genomes bearing *cyd–cox* regions with each other and with strains without *cyd–cox* regions (*P. caldoxylosilyticus* CIC9 for *Parageobacillus* genomosp. 1 NUB3621, and *Geobacillus* sp. E263 for *P. galactosidasius* DSM 18751). To investigate the cell functions conserved in genomes having *codhs*, presence or absence of orthologous genes identified above by Orthofinder was compared between genomes with and without the *coo–ech* gene cluster, and between genomes with *cyd–cox* region containing accessory genes of *cox*, and genomes without one.

### Cultivation in the presence of different terminal electron acceptors

G301 cells were cultured with and without CO, as follows: Frozen stock of G301 was stroked on the TGP agar plate (54) and incubated overnight at 65 °C under aerobic conditions. A single colony was injected into 5 mL of the modified B medium in a 180 mm×18 mm glass test tube (IWAKI) with a polycarbonate screw cap (IWAKI) and grown aerobically at 65 °C and pH 6.7. Growth was monitored by OD_600_, and when OD_600_ reached 0.49–0.56, which was around the exponential growth phase. Subsequently, 500 μL of the aliquot was injected into 50 mL of the modified B medium in 300 mL grass bottles (PYREX) sealed with bromobutyl rubber stoppers (Altair Corporation) and phenol resin screw caps (SIBATA) were used.

To characterize CO metabolism, G301 was cultured under three distinct conditions. G301 was cultured with neither O_2_ nor nitrate in the modified B medium described above under a 20% CO/80% N_2_ atmosphere or a 100% N_2_ atmosphere as a negative control at 65 °C and pH 6.8 with gyratory shaking at 100 rpm. Growth was monitored by measuring OD_600_. The gas composition was monitored as described above, and Ar was used as the carrier gas.

G301 was cultured under a 20% CO/80% N_2_ atmosphere or 100% N_2_ atmosphere as a negative control, as described above, with the exception of modified B medium including 5.0 g/L of KNO_3_. Growth and gas composition were monitored as described above. Nitrate and nitrite were quantified by Griess reaction using the NO_2_/NO_3_ Assay Kit CII (Dojindo Laboratories) according to the manufacturer’s protocol.

G301 was cultured under a 20% CO/80% or a 20% N_2_/80% air atmosphere as a negative control at 65 °C and pH 6.7 with gyratory shaking at 100 rpm. Growth and gas composition were monitored as described above, except for the gas vials purged with Ar to quantify O_2_ prior to analysis. Each of the three experiments above were conducted in triplicate. Significance of difference in growth yield and growth rate were analyzed by Student’s t-test.

In order to analyze whether *Parageobacillus* with only *cooS* can couple CO oxidation with nitrate reduction, we cultured *P. thermoglucosidasius* in 25% CO/75% N_2_ atmosphere in the presence and absence of nitrate (KNO_3_ or KCl was added at a final concentration of 50 mM, respectively) at 65 °C and pH 7.0. The medium was Bly medium, which had a similar composition to modified B medium but had the amount of yeast extract reduced by one-tenth.

## Data availability

The genome of strain G301 was deposited in GenBank under the name “BioProject PRJDB14871” (genome accession numbers BSDB01000001–01000009; BioSample SAMD00562293).

## Acknowledgements

This work was supported by JSPS KAKENHI Grant Number JP16H06381 (to Y. S.) and by the Institute for Fermentation, Osaka (to T.Y.). We would like to thank Editage (www.editage.com) for English language editing.

## Supplemental figures

**Fig. S1.**
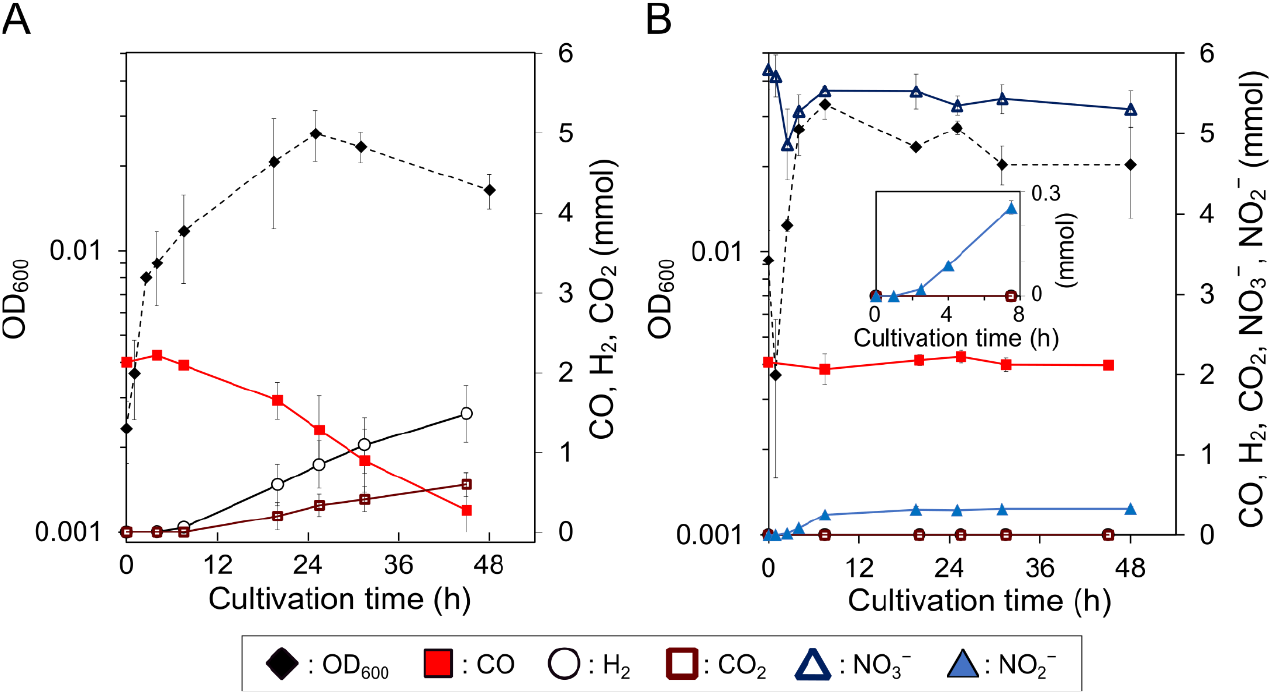
Growth of the *P. thermoglucosidasius* cultures and levels of CO, H_2_, CO_2_, nitrate, and nitrite. (A) OD_600_ and amount of CO, CO_2_, and H_2_ in the culture containing 50 mM KCl instead of KNO_3_. (B) OD_600_ and amount of CO, CO_2_, H_2_, nitrate, and nitrite in the culture containing 50 mM KNO_3_. The inlet graph shows the amount of nitrate and CO_2_ using a smaller scale. Plots represent the means of three biological replicates. Thin vertical lines represent standard deviations.

**Fig. S2.**
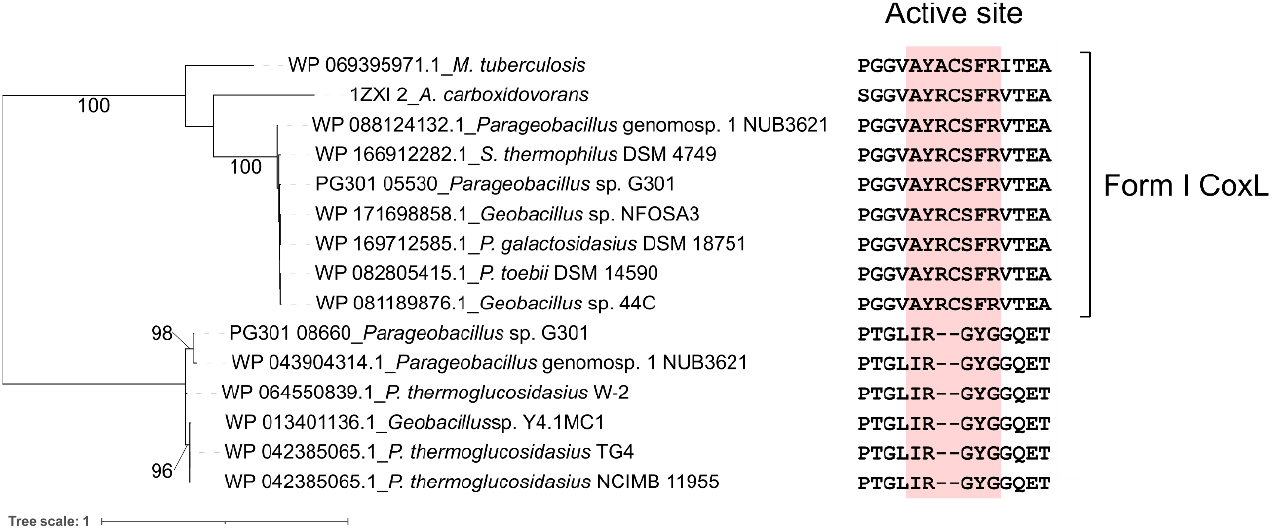
Maximum-likelihood tree of CoxL homologs and the active-site motifs. Bootstrap values >90% were shown on branches. The tree scale represents substitutions per site. CoxL of *M. tuberculosis* and *A. carboxidovorans* is known as Form I CoxL.

## References

1. Chin BY, Otterbein LE. 2009. Carbon monoxide is a poison … to microbes! CO as a bactericidal molecule. Curr Opin Pharmacol 9:490–500.

2. Coburn RF. 1979. Mechanisms of carbon monoxide toxicity. Prev Med 8:310–322.

3. Fukuyama Y, Inoue M, Omae K, Yoshida T, Sako Y. 2020. Chapter Three - Anaerobic and hydrogenogenic carbon monoxide-oxidizing prokaryotes: versatile microbial conversion of a toxic gas into an available energy, p. 99–148. *In* Gadd, GM, Sariaslani, S (eds.), Adv Appl Microbiol. Academic Press.

4. Thauer RK. 1990. Energy metabolism of methanogenic bacteria. Biochim Biophys Acta Bioenerg 1018:256–259.

5. King GM. 2006. Nitrate-dependent anaerobic carbon monoxide oxidation by aerobic CO-oxidizing bacteria. FEMS Microbiol Ecol 56:1–7.

6. King GM, Weber CF. 2007. Distribution, diversity and ecology of aerobic CO-oxidizing bacteria. Nat Rev Microbiol 5:107–118.

7. Myers M, King GM. 2017. Perchlorate-coupled carbon monoxide (CO) oxidation: evidence for a plausible microbe-mediated reaction in martian brines. Front Microbiol 8.

8. Robb FT, Techtmann SM. 2018. Life on the fringe: microbial adaptation to growth on carbon monoxide. F1000 Rep 7:F1000 Faculty Rev 1981.

9. Yoneda Y, Yoshida T, Kawaichi S, Daifuku T, Takabe K, Sako Y. 2012. *Carboxydothermus pertinax* sp. nov., a thermophilic, hydrogenogenic, Fe(III)-reducing, sulfur-reducing carboxydotrophic bacterium from an acidic hot spring. Int J Syst Evol Microbiol 62:1692–1697.

10. Slobodkin A, Slobodkina G, Allioux M, Alain K, Jebbar M, Shadrin V, Kublanov I, Toshchakov S, Bonch-Osmolovskaya E. 2019. Genomic insights into the carbon and energy metabolism of a thermophilic deep-sea bacterium *Deferribacter autotrophicus* revealed new metabolic traits in the phylum deferribacteres. 11. Genes 10:849.

11. Weghoff MC, Müller V. 2016. CO Metabolism in the thermophilic acetogen Thermoanaerobacter kivui. Appl Environ Microbiol 82:2312–2319.

12. Yoneda Y, Yoshida T, Yasuda H, Imada C, Sako Y. 2013. A thermophilic, hydrogenogenic and carboxydotrophic bacterium, *Calderihabitans maritimus* gen. nov., sp. nov., from a marine sediment core of an undersea caldera. Int J Syst Evol Microbiol 63:3602–3608.

13. Techtmann SM, Colman AS, Robb FT. 2009. ‘That which does not kill us only makes us stronger’: the role of carbon monoxide in thermophilic microbial consortia. Environ Microbiol 11:1027–1037.

14. Inoue M, Nakamoto I, Omae K, Oguro T, Ogata H, Yoshida T, Sako Y. 2019. Structural and phylogenetic diversity of anaerobic carbon-monoxide dehydrogenases. Front Microbiol 9.

15. Dobbek H, Gremer L, Meyer O, Huber R. 1999. Crystal structure and mechanism of CO dehydrogenase, a molybdo iron-sulfur flavoprotein containing S-selanylcysteine. Proc Natl Acad Sci U S A 96:8884–8889.

16. Arendsen AF, Hadden J, Card G, McAlpine AS, Bailey S, Zaitsev V, Duke EHM, Lindley PF, Kröckel M, Trautwein AX, Feiters MC, Charnock JM, Garner CD, Marritt SJ, Thomson AJ, Kooter IM, Johnson MK, van den Berg WAM, van Dongen WMAM, Hagen WR. 1998. The “prismane” protein resolved: X-ray structure at 1.7 Å and multiple spectroscopy of two novel 4Fe clusters. J Biol Inorg Chem 3:81–95.

17. Mistry J, Chuguransky S, Williams L, Qureshi M, Salazar GA, Sonnhammer ELL, Tosatto SCE, Paladin L, Raj S, Richardson LJ, Finn RD, Bateman A. 2021. Pfam: The protein families database in 2021. Nucleic Acids Res 49:D412–D419.

18. Techtmann SM, Lebedinsky AV, Colman AS, Sokolova TG, Woyke T, Goodwin L, Robb FT. 2012. Evidence for horizontal gene transfer of anaerobic carbon monoxide Dehydrogenases. Front Microbiol 3.

19. Inoue M, Omae K, Nakamoto I, Kamikawa R, Yoshida T, Sako Y. 2022. Biome-specific distribution of Ni-containing carbon monoxide dehydrogenases. Extremophiles 26.

20. Wang M, Tomb J-F, Ferry JG. 2011. Electron transport in acetate-grown Methanosarcina acetivorans. BMC Microbiol 11.

21. Alfano M, Cavazza C. 2018. The biologically mediated water–gas shift reaction: structure, function and biosynthesis of monofunctional [NiFe]-carbon monoxide dehydrogenases. Sustain Energy Fuels 2:1653–1670.

22. Hadj-Saïd J, Pandelia M-E, Léger C, Fourmond V, Dementin S. 2015. The carbon monoxide dehydrogenase from Desulfovibrio vulgaris. Biochim Biophys Acta Bioenerg 1847:1574–1583.

23. Dobbek H, Svetlitchnyi V, Gremer L, Huber R, Meyer O. 2001. Crystal structure of a carbon monoxide dehydrogenase reveals a [Ni-4Fe-5S] Cluster. Science 293:1281–1285.

24. Ensign SA, Ludden PW. 1991. Characterization of the CO oxidation/H_2_ evolution system of *Rhodospirillum rubrum*. Role of a 22-kDa iron-sulfur protein in mediating electron transfer between carbon monoxide dehydrogenase and hydrogenase. J Biol Chem 266:18395–18403.

25. Jeon WB, Cheng J, Ludden PW. 2001. Purification and characterization of membrane-associated CooC protein and its functional role in the insertion of nickel into carbon monoxide dehydrogenase from Rhodospirillum rubrum*. J Biol Chem 276:38602–38609.

26. Singer SW, Hirst MB, Ludden PW. 2006. CO-dependent H_2_ evolution by *Rhodospirillum rubrum*: Role of CODH:CooF complex. Biochim Biophys Acta Bioenerg 1757:1582–1591.

27. Schoelmerich MC, Müller V. 2019. Energy conservation by a hydrogenase-dependent chemiosmotic mechanism in an ancient metabolic pathway. Proc Natl Acad Sci U S A 116:6329–6334.

28. Schut GJ, Lipscomb GL, Nguyen DMN, Kelly RM, Adams MWW. 2016. Heterologous production of an energy-conserving carbon monoxide dehydrogenase complex in the hyperthermophile Pyrococcus furiosus. Front Microbiol 7.

29. Hille R. 1996. The mononuclear molybdenum enzymes. Chem Rev 96:2757–2816.

30. Maria. JR, Margarida A, Isabel M, Josee JGM, Jean L, Richard E, Monika S, Peter H, Robert H. 1995. Crystal structure of the xanthine oxidase-related aldehyde oxido-reductase from D. gigas. Science 270:1170–1176.

31. King GM. 2003. Molecular and culture-based analyses of aerobic carbon monoxide oxidizer diversity. Appl Environ Microbiol 69:7257–7265.

32. Choi ES, Min K, Kim G-J, Kwon I, Kim YH. 2017. Expression and characterization of *Pantoea* CO dehydrogenase to utilize CO-containing industrial waste gas for expanding the versatility of CO dehydrogenase. 1. Sci Rep 7:44323.

33. Kang BS, Kim YM. 1999. Cloning and molecular characterization of the genes for carbon monoxide dehydrogenase and localization of molybdopterin, flavin adenine dinucleotide, and iron-sulfur centers in the enzyme of Hydrogenophaga pseudoflava. J Bacteriol 181:5581–5590.

34. Wilcoxen J, Zhang B, Hille R. 2011. Reaction of the Molybdenum-and Copper-containing carbon monoxide dehydrogenase from *Oligotropha carboxydovorans* with quinones. Biochemistry 50:1910–1916.

35. Cordero PRF, Bayly K, Man Leung P, Huang C, Islam ZF, Schittenhelm RB, King GM, Greening C. 2019. Atmospheric carbon monoxide oxidation is a widespread mechanism supporting microbial survival. ISME J 13:2868–2881.

36. Sokolova TG, Yakimov MM, Chernyh NA, Lun’kova EYu, Kostrikina NA, Taranov EA, Lebedinskii AV, Bonch-Osmolovskaya EA. 2017. Aerobic carbon monoxide oxidation in the course of growth of a hyperthermophilic archaeon, *Sulfolobus* sp. ETSY. Microbiology 86:539–548.

37. Cypionka H, van Verseveld HW, Stouthamer AH. 1984. Proton translocation coupled to carbon monoxide-insensitive and -sensitive electron transport in *Pseudomonas carboxydovorans*. FEMS Microbiol Lett 22:209–213.

38. Nanba K, King GM, Dunfield K. 2004. Analysis of facultative lithotroph distribution and diversity on volcanic deposits by use of the large subunit of ribulose 1,5-bisphosphate carboxylase/oxygenase. Appl Environ Microbiol 70:2245–2253.

39. Santiago B, Schübel U, Egelseer C, Meyer O. 1999. Sequence analysis, characterization and CO-specific transcription of the cox gene cluster on the megaplasmid pHCG3 of *Oligotropha carboxidovorans*. Gene 236:115–124.

40. Lorite MJ, Tachil J, Sanjuán J, Meyer O, Bedmar EJ. 2000. Carbon monoxide dehydrogenase activity in Bradyrhizobium japonicum. Appl Environ Microbiol 66:1871–1876.

41. Hogendoorn C, Pol A, Picone N, Cremers G, van Alen TA, Gagliano AL, Jetten MSM, D’Alessandro W, Quatrini P, Op den Camp HJM. 2020. Hydrogen and carbon monoxide-utilizing *Kyrpidia spormannii* species from Pantelleria island, Italy. Front Microbiol 11.

42. Cunliffe M. 2011. Correlating carbon monoxide oxidation with *cox* genes in the abundant Marine Roseobacter Clade. ISME J 5:685–691.

43. Merrouch M, Hadj-Saïd J, Domnik L, Dobbek H, Léger C, Dementin S, Fourmond V. 2015. O_2_ inhibition of Ni-containing CO dehydrogenase is partly reversible. Chemistry 21:18934–18938.

44. Omae K, Fukuyama Y, Yasuda H, Mise K, Yoshida T, Sako Y. 2019. Diversity and distribution of thermophilic hydrogenogenic carboxydotrophs revealed by microbial community analysis in sediments from multiple hydrothermal environments in Japan. Arch Microbiol 201:969–982.

45. Artuso I, Turrini P, Pirolo M, Lucidi M, Tescari M, Visaggio D, Mansi A, Lugli GA, Ventura M, Visca P. 2021. Phylogenomic analysis and characterization of carbon monoxide utilization genes in the family Phyllobacteriaceae with reclassification of *Aminobacter carboxidus* (Meyer et al. 1993, Hördt et al. 2020) as *Aminobacter lissarensis* comb. nov. (McDonald et al. 2005). Syst Appl Microbiol 44:126199.

46. Coorevits A, Dinsdale AE, Halket G, Lebbe L, De Vos P, Van Landschoot A, Logan NA. 2012. Taxonomic revision of the genus *Geobacillus:* emendation of *Geobacillus, G. stearothermophilus, G*. *jurassicus*, *G. toebii*, *G. thermodenitrificans* and *G. thermoglucosidans* (nom. corrig., formerly ’*thermoglucosidasius*’); transfer of *Bacillus thermantarcticus* to the genus as G. thermantarcticus comb. nov.; proposal of *Caldibacillus* debilis gen. nov., comb. nov.; transfer of *G. tepidamans* to *Anoxybacillus* as *A. tepidamans* comb. nov.; and proposal of *Anoxybacillus caldiproteolyticus* sp. nov. Int J Syst Evol Microbiol 62:1470–1485.

47. Aliyu H, Lebre P, Blom J, Cowan D, De Maayer P. 2016. Phylogenomic re-assessment of the thermophilic genus Geobacillus. Syst Appl Microbiol 39:527–533.

48. Aliyu H, Lebre P, Blom J, Cowan D, De Maayer P. 2018. Corrigendum to “Phylogenomic re-assessment of the thermophilic genus *Geobacillus*” [Syst. Appl. Microbiol. 39 (2016) 527–533]. Syst Appl Microbiol 41:529–530.

49. Najar IN, Thakur N. 2020. A systematic review of the genera *Geobacillus* and *Parageobacillus:* their evolution, current taxonomic status and major applications. Microbiology 166:800–816.

50. Madhaiyan M, Saravanan VS, See-Too W-S. in press. Genome based analyses reveals the presence of heterotypic synonyms and subspecies in Bacteria and Archaea. bioRxiv https://doi.org/10.1101/2020.12.13.418756.

51. Saeed AM, El-Shatoury EH, Sayed HAE. 2021. Statistical factorial designs for optimum production of thermostable α-amylase by the degradative bacterium *Parageobacillus thermoglucosidasius* Pharon1 isolated from Sinai, Egypt. J Genet Eng Biotechnol 19.

52. Suleiman AD, Abdul Rahman N, Mohd Yusof H, Mohd Shariff F, Yasid NA. 2020. Effect of cultural conditions on protease production by a Thermophilic *Geobacillus thermoglucosidasius* SKF4 isolated from Sungai Klah hot spring park, Malaysia. Molecules 25:2609.

53. Bashir Z, Sheng L, Anil A, Lali A, Minton NP, Zhang Y. 2019. Engineering *Geobacillus thermoglucosidasius* for direct utilisation of holocellulose from wheat straw. Biotechnol Biofuels 12:199.

54. Cripps RE, Eley K, Leak DJ, Rudd B, Taylor M, Todd M, Boakes S, Martin S, Atkinson T. 2009. Metabolic engineering of *Geobacillus thermoglucosidasius* for high yield ethanol production. Metab Eng 11:398–408.

55. Yang Z, Sun Q, Tan G, Zhang Q, Wang Z, Li C, Qi F, Wang W, Zhang L, Li Z. 2021. Engineering thermophilic *Geobacillus thermoglucosidasius* for riboflavin production. Microb Biotechnol 14:363–373.

56. Mohr T, Aliyu H, Küchlin R, Polliack S, Zwick M, Neumann A, Cowan D, de Maayer P. 2018. CO-dependent hydrogen production by the facultative anaerobe *Parageobacillus thermoglucosidasius*. Microb Cell Fact 17.

57. Mohr T, Aliyu H, Küchlin R, Zwick M, Cowan D, Neumann A, de Maayer P. 2018. Comparative genomic analysis of *Parageobacillus thermoglucosidasius* strains with distinct hydrogenogenic capacities. BMC Genomics 19.

58. Inoue M, Tanimura A, Ogami Y, Hino T, Okunishi S, Maeda H, Yoshida T, Sako Y. 2019. Draft genome sequence of *Parageobacillus thermoglucosidasius* strain TG4, a hydrogenogenic carboxydotrophic bacterium isolated from a marine sediment. Microbiol Resour Announc 8.

59. Adachi Y, Inoue M, Yoshida T, Sako Y. 2021. Genetic engineering of carbon monoxide-dependent hydrogen-producing machinery in *Parageobacillus thermoglucosidasius*. Microbes Environ 35.

60. Wattam AR, Davis JJ, Assaf R, Boisvert S, Brettin T, Bun C, Conrad N, Dietrich EM, Disz T, Gabbard JL, Gerdes S, Henry CS, Kenyon RW, Machi D, Mao C, Nordberg EK, Olsen GJ, Murphy-Olson DE, Olson R, Overbeek R, Parrello B, Pusch GD, Shukla M, Vonstein V, Warren A, Xia F, Yoo H, Stevens RL. 2017. Improvements to PATRIC, the all-bacterial bioinformatics database and analysis resource center. Nucleic Acids Res 45:D535–D542.

61. Schoch CL, Ciufo S, Domrachev M, Hotton CL, Kannan S, Khovanskaya R, Leipe D, Mcveigh R, O’Neill K, Robbertse B, Sharma S, Soussov V, Sullivan JP, Sun L, Turner S, Karsch-Mizrachi I. 2020. NCBI Taxonomy: a comprehensive update on curation, resources and tools. Database 2020:baaa062.

62. Jain C, Rodriguez-R LM, Phillippy AM, Konstantinidis KT, Aluru S. 2018. High throughput ANI analysis of 90K prokaryotic genomes reveals clear species boundaries. 1. Nat Commun 9.

63. Søndergaard D, Pedersen CNS, Greening C. 2016. HydDB: a web tool for hydrogenase classification and analysis. Sci Rep 6:34212.

64. Winstedt L, Yoshida K-I, Fujita Y, von Wachenfeldt C. 1998. Cytochrome bd biosynthesis in *Bacillus subtilis:* characterization of the *cydABCD* Operon. J Bacteriol 180:6571–6580.

65. Magalon A, Fedor JG, Walburger A, Weiner JH. 2011. Molybdenum enzymes in bacteria and their maturation. Coord Chem Rev 255:1159–1178.

66. Sharma P, Teixeira de Mattos MJ, Hellingwerf KJ, Bekker M. 2012. On the function of the various quinone species in Escherichia coli. FEBS J 279:3364–3373.

67. Ragsdale SW. 2004. Life with carbon monoxide. Crit Rev Biochem Mol Biol 39:165–195.

68. Westphal L, Wiechmann A, Baker J, Minton NP, Müller V. 2018. The Rnf complex is an energy-coupled transhydrogenase essential to reversibly link cellular NADH and ferredoxin pools in the acetogen Acetobacterium woodii. J Bacteriol 200:e00357–18.

69. Soboh B, Linder D, Hedderich R. 2002. Purification and catalytic properties of a CO-oxidizing:H_2_-evolving enzyme complex from *Carboxydothermus hydrogenoformans*. European Journal of Biochemistry 269:5712–5721.

70. Wiechmann A, Trifunović D, Klein S, Müller V. 2020. Homologous production, one-step purification, and proof of Na^+^ transport by the Rnf complex from *Acetobacterium woodii*, a model for acetogenic conversion of C1 substrates to biofuels. Biotechnol Biofuels 13:208.

71. Ghachi ME, Bouhss A, Blanot D, Mengin-Lecreulx D. 2004. The *bacA* gene of *Escherichia coli* encodes an undecaprenyl pyrophosphate phosphatase activity *. J Biol Chem 279:30106–30113.

72. Chiodini G, Caliro S, Caramanna G, Granieri D, Minopoli C, Moretti R, Perotta L, Ventura G. 2006. Geochemistry of the submarine gaseous emissions of Panarea (Aeolian Islands, Southern Italy): magmatic vs. hydrothermal origin and implications for volcanic surveillance. Pure appl geophys 163:759–780.

73. Schade GW, Crutzen PJ. 1999. CO emissions from degrading plant matter (II). Tellus B Chem Phys Meteorol 51:909–918.

74. Conte L, Szopa S, Séférian R, Bopp L. 2019. The oceanic cycle of carbon monoxide and its emissions to the atmosphere. Biogeosciences 16:881–902.

75. Zuo Y, Jones RD. 1997. Photochemistry of natural dissolved organic matter in lake and wetland waters—production of carbon monoxide. Water Res 31:850–858.

76. Montag D, Schink B. 2018. Formate and hydrogen as electron shuttles in terminal fermentations in an oligotrophic freshwater lake sediment. Appl Environ Microbiol 84:e01572–18.

77. Cunliffe M. 2013. Physiological and metabolic effects of carbon monoxide oxidation in the model marine bacterioplankton *Ruegeria pomeroyi* DSS-3. Appl Environ Microbiol 79.

78. Balk M, Heilig HGHJ, van Eekert MHA, Stams AJM, Rijpstra IC, Sinninghe-Damsté JS, de Vos WM, Kengen SWM. 2009. Isolation and characterization of a new CO-utilizing strain, *Thermoanaerobacter thermohydrosulfuricus* subsp. carboxydovorans, isolated from a geothermal spring in Turkey. Extremophiles 13:885–894.

79. Jain S, Katsyv A, Basen M, Müller V. 2021. The monofunctional CO dehydrogenase CooS is essential for growth of *Thermoanaerobacter kivui* on carbon monoxide. Extremophiles 26:4.

80. Pfennig N. 1974. *Rhodopseudomonas globiformis*, sp. n., a new species of the Rhodospirillaceae. Arch Microbiol 100:197–206.

81. Wolin EA, Wolin MJ, Wolfe RS. 1963. Formation of methane by bacterial extracts. J Biol Chem 238:2882–2886.

82. Yamamura H, Hayashi T, Hamada M, Kohda T, Serisawa Y, Matsuyama-Serisawa K, Nakagawa Y, Otoguro M, Yanagida F, Tamura T, Hayakawa M. 2019. *Cellulomonas algicola* sp. nov., an actinobacterium isolated from a freshwater alga. Int J Syst Evol Microbiol 69:2723–2728.

83. Fukuyama Y, Tanimura A, Inoue M, Omae K, Yoshida T, Sako Y. 2019. Draft genome sequences of two thermophilic *Moorella* sp. strains, isolated from an acidic hot spring in Japan. Microbiol Resour Announc 8:e00663–19.

84. Watson ML. 1958. Staining of tissue sections for electron microscopy with heavy metals. J Biophys Biochem Cytol 4:475–478.

85. Kurokawa M, Nakano M, Kitahata N, Kuchitsu K, Furuya T. 2021. An efficient direct screening system for microorganisms that activate plant immune responses based on plant–microbe interactions using cultured plant cells. Sci Rep 11:7396.

86. Bolger AM, Lohse M, Usadel B. 2014. Trimmomatic: a flexible trimmer for Illumina sequence data. Bioinformatics 30:2114–2120.

87. O’Connell J, Schulz-Trieglaff O, Carlson E, Hims MM, Gormley NA, Cox AJ. 2015. NxTrim: optimized trimming of Illumina mate pair reads. Bioinformatics 31:2035–2037.

88. Bankevich A, Nurk S, Antipov D, Gurevich AA, Dvorkin M, Kulikov AS, Lesin VM, Nikolenko SI, Pham S, Prjibelski AD, Pyshkin AV, Sirotkin AV, Vyahhi N, Tesler G, Alekseyev MA, Pevzner PA. 2012. SPAdes: a new genome assembly algorithm and its applications to single-cell sequencing. J Comput Biol 19:455–477.

89. Li H, Durbin R. 2010. Fast and accurate long-read alignment with Burrows-Wheeler transform. Bioinformatics 26:589–595.

90. Li H, Handsaker B, Wysoker A, Fennell T, Ruan J, Homer N, Marth G, Abecasis G, Durbin R, 1000 genome project data processing subgroup. 2009. the sequence alignment/map format and SAMtools. Bioinformatics 25:2078–2079.

91. Murphy RR, O’Connell J, Cox AJ, Schulz-Trieglaff O. 2015. NxRepair: error correction in de novo sequence assembly using Nextera mate pairs. PeerJ 3:e996.

92. Tanizawa Y, Fujisawa T, Nakamura Y. 2018. DFAST: a flexible prokaryotic genome annotation pipeline for faster genome publication. Bioinformatics 34:1037–1039.

93. O’Leary NA, Wright MW, Brister JR, Ciufo S, Haddad D, McVeigh R, Rajput B, Robbertse B, Smith-White B, Ako-Adjei D, Astashyn A, Badretdin A, Bao Y, Blinkova O, Brover V, Chetvernin V, Choi J, Cox E, Ermolaeva O, Farrell CM, Goldfarb T, Gupta T, Haft D, Hatcher E, Hlavina W, Joardar VS, Kodali VK, Li W, Maglott D, Masterson P, McGarvey KM, Murphy MR, O’Neill K, Pujar S, Rangwala SH, Rausch D, Riddick LD, Schoch C, Shkeda A, Storz SS, Sun H, Thibaud-Nissen F, Tolstoy I, Tully RE, Vatsan AR, Wallin C, Webb D, Wu W, Landrum MJ, Kimchi A, Tatusova T, DiCuccio M, Kitts P, Murphy TD, Pruitt KD. 2016. Reference sequence (RefSeq) database at NCBI: current status, taxonomic expansion, and functional annotation. Nucleic Acids Res 44:D733–D745.

94. Parks DH, Chuvochina M, Rinke C, Mussig AJ, Chaumeil P-A, Hugenholtz P. 2022. GTDB: an ongoing census of bacterial and archaeal diversity through a phylogenetically consistent, rank normalized and complete genome-based taxonomy. Nucleic Acids Res 50:D785–D794.

95. Gurevich A, Saveliev V, Vyahhi N, Tesler G. 2013. QUAST: quality assessment tool for genome assemblies. Bioinformatics 29:1072–1075.

96. Katoh K, Kuma K, Toh H, Miyata T. 2005. MAFFT version 5: improvement in accuracy of multiple sequence alignment. Nucleic Acids Res 33:511–518.

97. Capella-Gutiérrez S, Silla-Martínez JM, Gabaldón T. 2009. trimAl: a tool for automated alignment trimming in large-scale phylogenetic analyses. Bioinformatics 25:1972–1973.

98. Nguyen L-T, Schmidt HA, von Haeseler A, Minh BQ. 2015. IQ-TREE: a fast and effective stochastic algorithm for estimating maximum-likelihood phylogenies. Mol Biol Evol 32:268–274.

99. Ivica L, Peer B. 2019. Interactive Tree Of Life (iTOL) v4: recent updates and new developments. Nucleic Acids Res 47:W256–W259.

100. Emms DM, Kelly S. 2019. OrthoFinder: phylogenetic orthology inference for comparative genomics. Genome Biol 20:238.

101. Edgar RC. 2004. MUSCLE: multiple sequence alignment with high accuracy and high throughput. Nucleic Acids Res 32:1792–1797.

102. Sudhir K, Glen S, Michael L, Christina K, Koichiro T. 2018. MEGA X: Molecular Evolutionary genetics analysis across computing platforms. Mol Biol Evol 35:1547–1549.

103. Meyer O, Schlegel HG. 1978. Reisolation of the carbon monoxide utilizing hydrogen bacterium *Pseudomonas carboxydovorans* (Kistner) comb. nov. Arch Microbiol 118:35–43.

104. Park SW, Hwang EH, Park H, Kim JA, Heo J, Lee KH, Song T, Kim E, Ro YT, Kim SW, Kim YM. 2003. Growth of Mycobacteria on carbon monoxide and methanol. J Bacteriol 185:142–147.

105. Buchfink B, Xie C, Huson DH. 2015. Fast and sensitive protein alignment using DIAMOND. 1. Nat Methods 12:59–60.

106. Delcher AL, Kasif S, Fleischmann RD, Peterson J, White O, Salzberg SL. 1999. Alignment of whole genomes. Nucleic Acids Res 27:2369–2376.

107. Nishimura Y, Yoshida T, Kuronishi M, Uehara H, Ogata H, Goto S. 2017. ViPTree: the viral proteomic tree server. Bioinformatics 33:2379–2380.

108. Singh NK, Carlson C, Sani RK, Venkateswaran K. 2017. Draft genome sequences of thermophiles isolated from Yates shaft, a deep-subsurface environment. Genome Announc 5:e00405–17.

109. Hatmaker EA, O’Dell KB, Riley LA, Payne IC, Guss AM. 2020. Methylome and complete genome sequence of *Parageobacillus toebii* DSM 14590^T^, a thermophilic bacterium. Microbiol Resour Announc 9:e00589–20.

110. Talamantes-Becerra B, Carling J, Kilian A, Georges A. 2020. Discovery of thermophilic Bacillales using reduced-representation genotyping for identification. BMC Microbiol 20.

111. Ramaloko WT, Koen N, Polliack S, Aliyu H, Lebre PH, Mohr T, Oswald F, Zwick M, Zeigler DR, Neumann A, Syldatk C, Cowan DA, De Maayer P. 2018. High quality draft genomes of the type strains *Geobacillus thermocatenulatus* DSM 730^T^, *G. uzenensis* DSM 23175^T^ and *Parageobacillus galactosidasius* DSM 18751^T^. J Genomics 6:20–23.

112. Sharma P, Gupta S, Sourirajan A, Baumler DJ, Dev K. 2019. Draft genome sequence of hyperthermophilic, halotolerant *Parageobacillus toebii* PW12, isolated from the Tattapani hot spring, northwest himalayas. Microbiol Resour Announc 8:e01163–18.

113. Najar IN, Sherpa MT, Das S, Thakur N. 2018. Draft genome sequence of *Geobacillus yumthangensis* AYN2 sp. nov., a denitrifying and sulfur reducing thermophilic bacterium isolated from the hot springs of Sikkim. Gene Rep 10:162–166.

114. Najar IN, Sherpa MT, Das S, Das S, Thakur N. 2020. Diversity analysis and metagenomic insights into antibiotic and metal resistance among Himalayan hot spring bacteriobiome insinuating inherent environmental baseline levels of antibiotic and metal tolerance. J Glob Antimicrob Resist 21:342–352.

115. Berendsen EM, Wells-Bennik MHJ, Krawczyk AO, de Jong A, van Heel A, Holsappel S, Eijlander RT, Kuipers OP. 2016. Draft genome sequences of seven thermophilic spore-forming bacteria isolated from foods that produce highly heat-resistant spores, comprising *Geobacillus* spp., *Caldibacillus debilis*, and *Anoxybacillus flavithermus*. Genome Announcements 4:e00105–16.

116. Chen Y, Wei D, Wang Y, Zhang X. 2013. The role of interactions between bacterial chaperone, aspartate aminotransferase, and viral protein during virus infection in high temperature environment: the interactions between bacterium and virus proteins. BMC Microbiol 13.

117. Brumm PJ, Land ML, Mead DA. 2016. Complete genome sequences of *Geobacillus* sp. WCH70, a thermophilic strain isolated from wood compost. Stand Genomic Sci 11:33.

118. Zhu L, Li M, Guo S, Wang W. 2016. Draft genome sequence of a thermophilic desulfurization bacterium, *Geobacillus thermoglucosidasius* strain W-2. Genome Announc 4:e00793–16.

119. Blanchard K, Robic S, Matsumura I. 2014. Transformable facultative thermophile *Geobacillus stearothermophilus* NUB3621 as a host strain for metabolic engineering. Appl Microbiol Biotechnol 98:6715–6723.

120. Lama L, Calandrelli V, Gambacorta A, Nicolaus B. 2004. Purification and characterization of thermostable xylanase and beta-xylosidase by the thermophilic bacterium *Bacillus thermantarcticus*. Res Microbiol 155:283–289.

121. Brumm PJ, Land ML, Mead DA. 2015. Complete genome sequence of *Geobacillus thermoglucosidasius* C56-YS93, a novel biomass degrader isolated from obsidian hot spring in Yellowstone National Park. Stand Genomic Sci 10:73.

122. Brumm P, Land ML, Hauser LJ, Jeffries CD, Chang Y-J, Mead DA. 2015. Complete genome sequence of *Geobacillus* strain Y4.1MC1, a novel CO-utilizing *Geobacillus thermoglucosidasius* strain isolated from Bath hot spring in Yellowstone National Park. Bioenerg Res 8:1039–1045.

123. Valladares Juárez AG, Dreyer J, Göpel PK, Koschke N, Frank D, Märkl H, Müller R. 2009. Characterisation of a new thermoalkaliphilic bacterium for the production of high-quality hemp fibres, *Geobacillus thermoglucosidasius* strain PB94A. Appl Microbiol Biotechnol 83:521–527.

124. Sheng L, Zhang Y, Minton NP. 2016. Complete genome sequence of *Geobacillus thermoglucosidasius* NCIMB 11955, the progenitor of a bioethanol production strain. Genome Announc 4:e01065–16.

125. Zhao Y, Caspers MP, Abee T, Siezen RJ, Kort R. 2012. Complete genome sequence of *Geobacillus thermoglucosidans* TNO-09.020, a thermophilic sporeformer associated with a dairy-processing environment. J Bacteriol 194:4118–4118.

126. Ahmad S, Scopes RK, Rees GN, Patel BK. 2000. *Saccharococcus caldoxylosilyticus* sp. nov., an obligately thermophilic, xylose-utilizing, endospore-forming bacterium. Int J Syst Evol Microbiol 50:517–523.

127. Studholme DJ. 2015. Some (bacilli) like it hot: genomics of *Geobacillus* species. Microb Biotechnol 8:40–48.

128. Imaura Y, Okamoto S, Hino T, Ogami Y, Katayama AY, Tanimura A, Inoue M, Kamikawa R, Yoshida T, Sako Y. 2023. The draft genome of G301. Accession numbers: BSDB01000001–01000009.

